# Replication Protein A recognizes DNA junctions to spatially organize genome maintenance

**DOI:** 10.64898/2026.11.24.769028

**Authors:** Ayush Mistry, Rahul Chadda, Ethan Hasenoehrl, Aidan Antony, Mansha Jain, Ishika Katukam, Tania Sultana, Monica Tokmina-Lukaszewska, Brian Bothner, Edwin Antony

## Abstract

Single-stranded DNA (ssDNA) intermediates arise during many DNA metabolic processes^1–3^ and are rapidly coated by the conserved ssDNA-binding protein Replication Protein A (RPA), which protects ssDNA and coordinates the recruitment of downstream DNA metabolic factors^4–7^. However, how RPA directs the precise spatial recruitment of DNA metabolic proteins to RPA-coated DNA remains unclear. Here we show that RPA possesses an intrinsic preference for single-stranded/double-stranded DNA (ss/dsDNA) junctions, creating localized binding states that promote the junction-specific recruitment of incoming proteins. Junction-proximal RPA exhibits enhanced resistance to disassembly through conserved structural elements within its DNA-binding domains. RPA phosphorylation diminishes this junction preference, providing a mechanism for remodeling junction-associated complexes. We further demonstrate that RPA directly regulates the junction-specific loading and engagement of the nucleotide excision repair factor XPA. Together, these findings establish a mechanistic basis for RPA recognition of ss/dsDNA junctions and support a general framework in which RPA spatially organizes DNA metabolic factors at sites of ssDNA formation. More broadly, these findings resolve how proteins gain selective access to DNA embedded within RPA-coated intermediates and identify junction recognition as a previously unrecognized regulatory feature of RPA function.

## Introduction

Replication Protein A (RPA) is an essential single-stranded DNA (ssDNA) binding protein that coordinates most DNA metabolic processes, including DNA replication, recombination, and repair^4–7^. RPA is a constitutive heterotrimer, comprised of the RPA70, RPA32, and RPA14 subunits. Its six oligonucleotide/oligosaccharide-binding (OB) folds are functionally classified as four DNA-binding domains (DBDs; A, B, C, and D) and two protein-interacting domains (PIDs; PID^70N^, and PID^32C^) (**Supplementary Fig. 1a**), allowing it to bind ssDNA with high affinity, and function as a hub to interact with and recruit over three dozen DNA metabolic proteins onto DNA^8^. Disordered linkers connect these domains and impart conformational flexibility to the complex^9–12^.

Despite its broad functional repertoire, RPA is frequently viewed primarily as a passive protector of ssDNA, and prevailing models of RPA function have largely developed in the context of ssDNA binding alone. However, many DNA metabolic processes occur at ss/dsDNA junctions, where newly exposed ssDNA is rapidly occupied by RPA. Notably, proteins involved in diverse DNA metabolic pathways preferentially engage these junctions. DNA damage checkpoint factors, including the human 9-1-1 clamp, the alternative yeast Rad24-RFC complex, and yeast Dpb11, localize to ss/dsDNA junctions during checkpoint activation^13,14^. Similarly, DNA repair proteins such as XPA during nucleotide excision repair, Rad52-Rad51 during homologous recombination, and the Smc5/6 complex during replication fork restart exhibit preferential engagement and/or prolonged residence at DNA junctions^15–17^. Recent studies further show that the microhomology-dependent end joining (MMEJ) factor Polθ initially engages short ssDNA overhangs at DNA junctions as a critical step in coordinating repair^18,19^. Collectively, these observations reveal a recurring but unexplained feature of DNA metabolism: proteins from diverse pathways preferentially operate at ss/dsDNA junctions that are typically occluded by RPA. Although RPA is ideally positioned to provide spatial information at these sites, whether it possesses intrinsic mechanisms to recognize DNA junctions and thereby direct the recruitment and activity of downstream factors remains unknown.

In this study, we address this question and establish a mechanistic framework for how RPA recognizes and responds to ss/dsDNA junctions. We show that RPA exhibits distinct interaction dynamics at DNA junctions and identify conserved features within its DNA-binding domains that regulate junction-specific engagement. We further demonstrate that phosphorylation of RPA modulates these interactions, providing a mechanism to control junction occupancy and junction-specific handoff of RPA-bound ssDNA to incoming proteins. As a proof of concept, we show that RPA directly regulates junction-specific recruitment of XPA. Our findings support a model in which RPA senses DNA architecture to encode positional information at ss/dsDNA junctions, thereby directing the localized recruitment and activity of downstream DNA metabolic factors. These results redefine the canonical view of RPA, from a passive ssDNA-binding factor to an active interpreter of DNA structure and organizer of genome maintenance.

## Results

### RPA molecules display preferential engagement to junction-proximal regions

RPA is a well-established, high-affinity ssDNA binding protein, but whether it possesses any preference for junction-proximal regions, especially when placed within the context of long ssDNA stretches is not known. To test this, we used a correlative optical tweezers fluorescence microscopy (CTFM) setup involving two optically-trapped polystyrene beads (∼4.0-4.9 µm in diameter). A precursor DNA substrate was tethered to the beads and subsequently manipulated using force to generate a gapped template containing both 5′ and 3′ junctions (total length of 17853 bp, with a 5005-nt ssDNA gap; **Fig. 1a, b**). We performed facilitated exchange experiments with RPA using two protein imaging channels, where site-specific fluorescently labeled RPA (RPA^MB543^; **Supplementary Fig. 1b**) was flown through the first protein imaging channel, and the second imaging channel carried excess, unlabeled RPA (**Fig. 1b, c**). Consistent with the exchange of DNA-bound RPA, we observed a loss of RPA^MB543^ in the second imaging channel (**Fig. 1c; Supplementary Fig. 2a**). However, we observed that while internally-bound RPA molecules on the 5005-nt ssDNA region displayed rapid exchange dynamics, junction-proximal RPA molecules resisted exchange for significantly longer periods of time before being evicted (**Fig. 1d**). This behavior prompted us to ask whether RPA displays any distinct engagement preferences for the junction-proximal regions. To test this, we performed continuous image scans of the entire DNA substrate as it entered the protein imaging channel with RPA^MB543^ (**Fig. 1e**). We found that RPA preferentially initiated engagement in the junction-proximal regions (∼1-5 s), which was promptly followed by RPA decorating the entire 5005-nt ssDNA gap (∼30 s; **Fig. 1e, f; Supplementary Fig. 2b**). An ATTO 647N dye was positioned on the dsDNA region 177 bp away from the 3ʹ recessed allowing us to identify the junctions. RPA displayed no inter-junction preference, with engagement initiating comparably at both the 5ʹ and 3ʹ junction-proximal regions (**Fig. 1f**). These findings reveal that in the context of gaps, or overhang-containing transient ssDNA intermediates, RPA preferentially engages the junction-proximal region and displays remarkably high intrinsic stability. These junction-specific engagement properties were also consistently observed with yeast Rpa (*Saccharomyces cerevisiae*) (**Supplementary Fig 3a, b**).

**Fig. 1.**
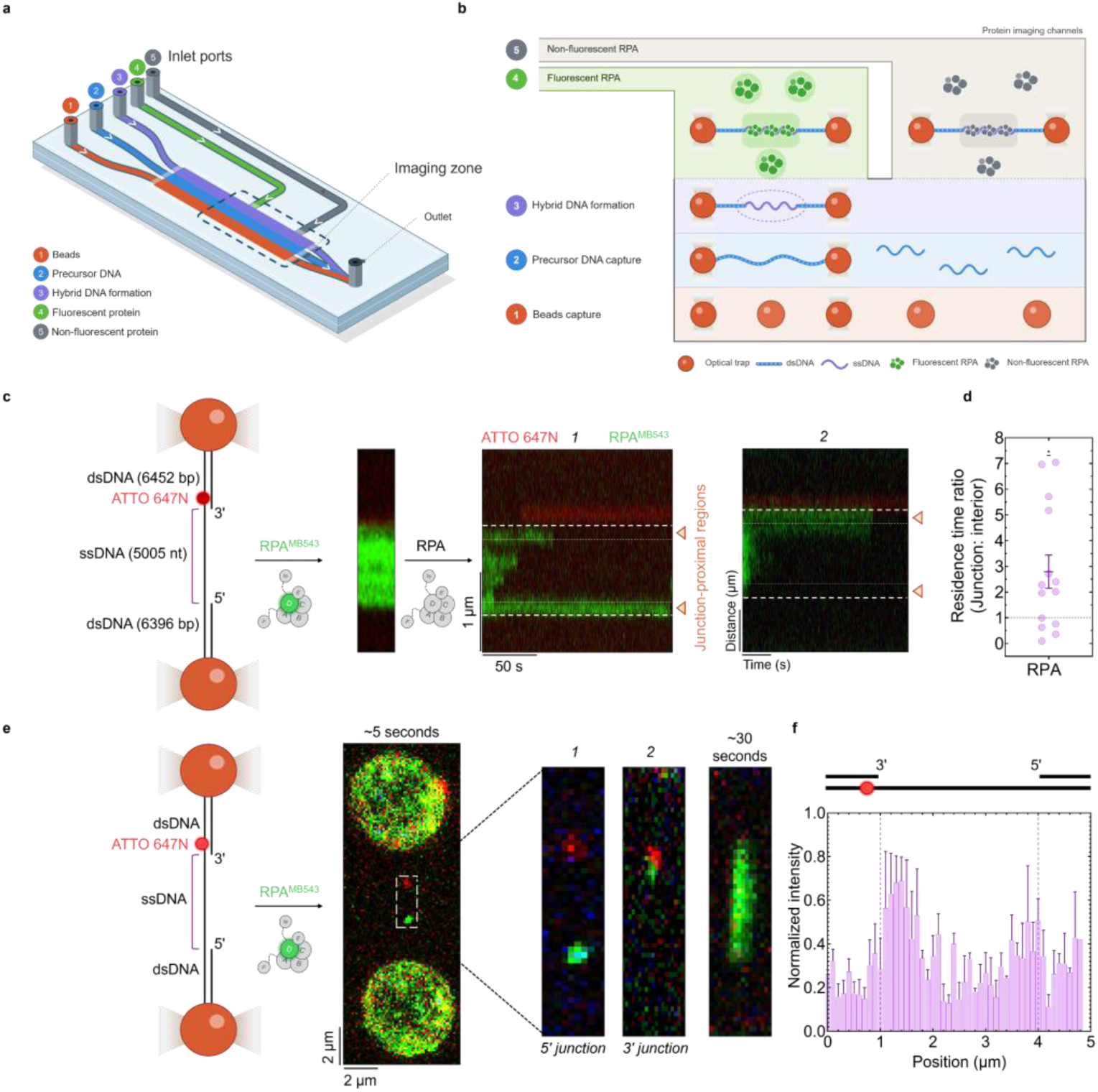
RPA preferentially and stably interacts with ss/dsDNA junctions. **a**, The flow-cell configuration of the single-molecule imaging setup. **b**, The experimental workflow of the single-molecule imaging assay. **c**, Optical trap assay to probe facilitated exchange on gapped DNA, where RPA^MB543^ (0.25-0.5 nM) was introduced in channel 4, allowed to coat the entire ssDNA gap, then challenged with excess unlabeled RPA (2.5 – 5 nM) in channel 5. The white dashed and dotted lines denote the area used to define and quantify ss/dsDNA junction & proximal regions in the kymograph. Atto647N at the 3ʹ junction marks junction polarity. Two representative kymographs are shown to depict 5ʹ (example *1*) and 3ʹ (example *2*) junction-proximal resistance to exchange. **d**, Junction-proximal: internal RPA^MB543^ residence time ratios plotted relative to a baseline ratio of 1 (1 indicates equivalent residence times); mean ± SEM is depicted, each data point is a single junction: internal residence time ratio. Data collected from N= 14 binding events. The non-parametric one-sample Wilcoxon test was applied to compare the median with a fixed median value of 1. (*) indicates p-value ≤ 0.033. **e**, Optical trap assay to capture initial engagement of RPA^MB543^, where RPA^MB543^ (0.25-0.5 nM) was introduced in channel 4, and the entire frame was continuously scanned. The first frame with detectable binding is depicted as a representative case. Two exemplar instances are provided to depict both 5ʹ (example *1*) and 3ʹ (example 2) junction-proximal initial engagement. **f**, Bar graph depicting the normalized MB543 fluorescence intensity as a function of position along the gapped DNA substrate. Light dotted lines highlight the ss/dsDNA junction. Mean intensity ± SEM plotted.

### Single-molecule FRET reveals differential engagement of RPA’s OB-folds on junction-containing DNA substrates

How RPA acquires this preference for DNA junctions remains an open question. We reasoned that this property may emerge from the differential engagement of its constituent DNA-binding and protein-interaction domains. Consistent with this idea, our previous work demonstrated that RPA reorganizes into distinct configurational states, defined by changes in the relative positioning of individual domains, in response to protein interactions and post-translational modifications (PTMs)^20–23^. We therefore considered whether DNA junction engagement similarly alters the configurational landscape of RPA. Given the distinct macroscopic modes by which RPA engages junction-containing and uninterrupted ssDNA substrates, we asked whether its DNA-binding domains display different patterns of engagement at a DNA junction relative to full ssDNA^7,24,25^.

Using the non-canonical amino acid (ncAA) incorporation strategy, we site-specifically engineered two RPA constructs to carry a fluorophore (RPA^Cy3^) on either RPA^DBD-A^ or RPA^DBD-D26^ (**Supplementary Fig. 1b**). By positioning the other fluorophore (Cy5) at either the DNA overhang end, or at the junction, we were able to use single-molecule Förster Resonance Energy Transfer (smFRET) to capture the dynamics of RPA’s DBDs as it sampled the junction-containing substrates (**Fig. 2**). Biotinylated DNA were tethered onto PEG-streptavidin coated glass slides and fluorescence changes in both the Cy3 and Cy5 channels were measured using single-molecule total internal reflectance microscopy (smTIRF). The site-size of RPA on ssDNA is ∼18-22 nt^27^, and a 25-mer ssDNA is canonically used to ensure that all the domains of RPA have sufficient space to engage. Here, we used a shorter (dT)_15_ overhang to promote configurational rearrangements of the domains, which could help us distinguish potential junction-specific effects (**Supplementary Table 1**).

**Fig. 2.**
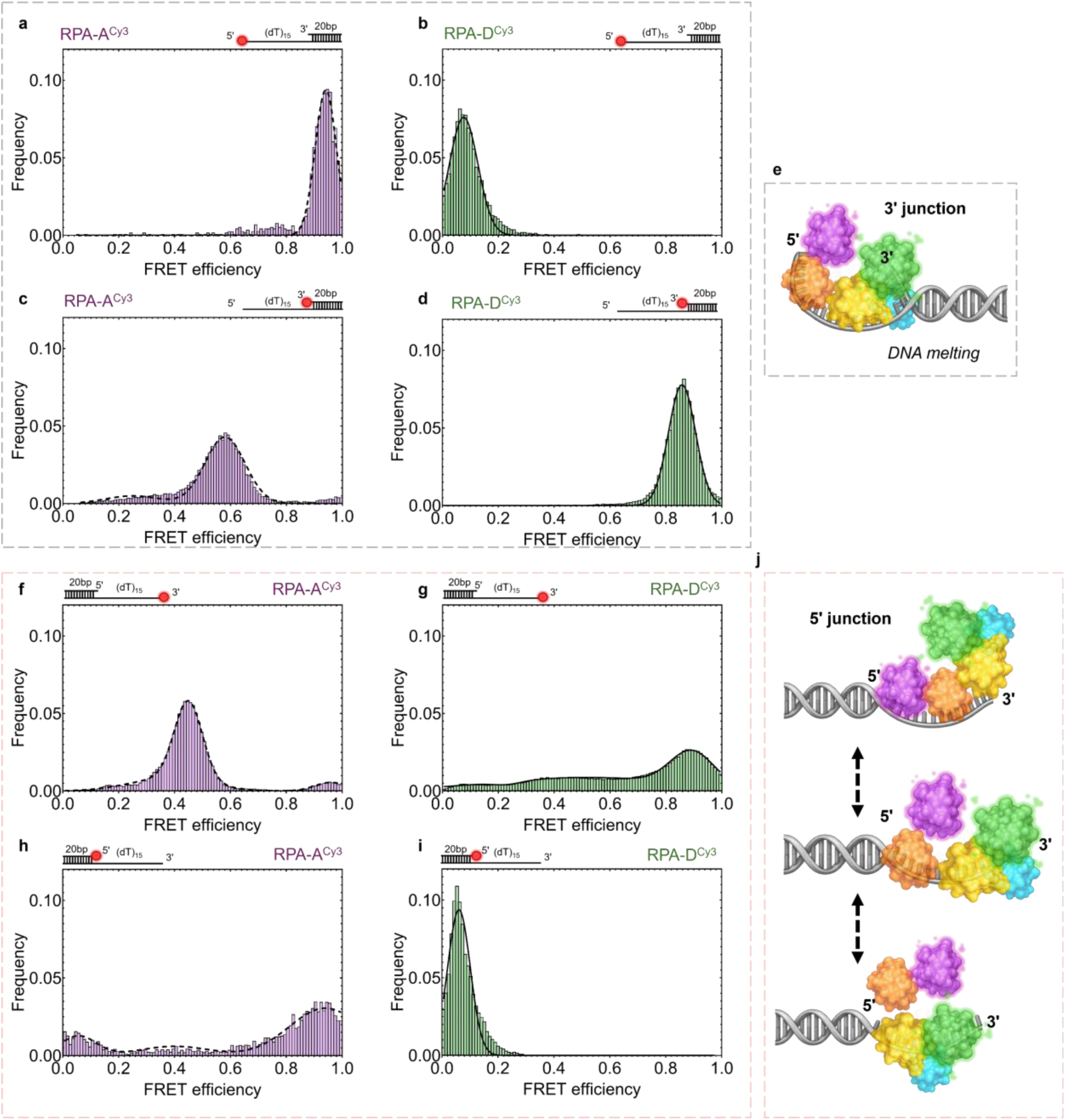
RPA distinctly samples DNA overhangs in a junction-specific manner. **a-h, f-i**, smFRET efficiencies depict the spatial positions of RPA DBD-A^Cy3^ or DBD-D^Cy3^ relative to end-labeled or junction-labeled Cy5 acceptor on the respective (dT)_15_ overhang DNA substrates. The models represent hypothetical configurations of the four DNA binding domains of RPA (DBD-A, magenta; DBD-B, orange; DBD-C, yellow; DBD-D, green; and OB-E, cyan (RPA14) on either the **e**, 3ʹ junction, or **j**, 5ʹ junction. The models are drawn to satisfy all the smFRET signatures for each junction. A single stable configuration bound to a melted 3ʹ junction is depicted. Multiple configurations are needed to fit the smFRET distributions on the 5ʹ junction.

On substrates with a 3ʹ junction and a Cy5 positioned at the terminus of the 5ʹ overhang, RPA^DBD-A-Cy3^ produced a predominantly high-FRET state, while RPA^DBD-D-Cy3^ generated a low-FRET signal (**Fig. 2a, b**). These FRET states are consistent with the expected polarity of RPA’s DBDs in engaging the ssDNA region, where DBD-A orients itself towards the 5ʹ end, and DBD-D towards the 3ʹ end^12,28,29^. When the Cy5 fluorophore was positioned at the 3ʹ junction, a mid-FRET state for RPA^DBD-A-Cy3^ (**Fig. 2c**), and a high-FRET state for RPA^DBD-D-Cy3^ (**Fig. 2d**), were observed, respectively. An integrative model that satisfies all four smFRET states requires ssDNA wrapping around the DBDs as observed in structural studies^20,30^, and partial melting/breathing of the duplex DNA (**Fig. 2e**). In support of this model, a 5′->3′ directional diffusion-based melting of dsDNA has been shown for both human and yeast RPA^27,31,32^. The data also demonstrates that RPA at the 3′ junction adopts a stable configuration.

Similar experiments carried out on a substrate with a 5′ junction and a (dT)_15_ overhang show a more dynamic smFRET landscape (**Fig. 2f-i**). When the Cy5 acceptor is positioned at the 3′ overhang, RPA^DBD-A-Cy3^ produced a major mid-FRET state (**Fig. 2f**), whereas RPA^DBD-D-Cy3^ generated a very broad distribution of states that spanned the entire spectrum of FRET efficiency, with a slight propensity for a high-FRET state (**Fig. 2g**). In corollary experiments where the Cy5 was moved to the 5′ junction, RPA^DBD-A-Cy3^ again produced a broad distribution of FRET states whereas RPA^DBD-D-Cy3^ displayed a single low FRET state. The first conclusion is that DBD-A (and likely DBD-B) is more dynamic at the 5′ junction. These data agree with the high-dynamic states that we previously showcased for DBDs-A & B^21^. Additional support for this model arises from smFRET analysis of a truncated version of RPA containing only domains F, A, & B where only transient FRET traces are captured compared to longer-lived states for the full-length proteins (**Supplementary Fig. 4**). The second conclusion is that the stability at the 5′ junction primarily arises from stable interactions of the trimerization core (Tri-C) of RPA. The data on the 5ʹ junction suggest a model that relies on the dynamic remodeling of RPA (**Fig. 2j**).

### Structural mass-spectrometry reveals minimal changes induced by the junction-containing substrates

Given the differences in the dynamics of RPA at the 5ʹ and 3ʹ junctions, we used hydrogen-deuterium exchange mass-spectrometry (HDX-MS) to elucidate any structure-based contributions. The four DNA-binding domains (DBD-A, B, C, and D) of RPA engage ssDNA and do not interact with dsDNA. The protein interaction domains (PID^70N^ and PID^32C^) are not known to interact with ssDNA. In HDX-MS experiments, we measured the uptake or loss of deuterium in response to DNA binding as a function of time. We have previously shown ssDNA-binding induced changes in the DBDs of RPA using HDX-MS^22^, and that ssDNA passively wraps around a C-shaped organization of the DBDs^20^. Here, we performed HDX-MS analysis of human RPA in the absence or presence of junction-containing DNA substrates. The substrates were similar to those used in the smFRET studies, with a 20-bp dsDNA region, and either 3ʹ or 5ʹ (dT)_15_ ssDNA overhangs (the complete peptide coverage is provided in **Supplementary Figures 5, 6, and 7**). To isolate junction-specific differences, we compared the HDX-MS profiles of unbound, 3ʹ, and 5ʹ bound RPA after 3 minutes of incubation and mapped these changes onto an AlphaFold3 model (**Fig. 3a**). While junction binding reduced exchange, there were no significant differences between the two junction-bound conditions, with a less than 5% deuterium uptake/loss overall (**Fig. 3a**). In a very minor subset of regions, we observe subtle differences across the RPA subunits, primarily in RPA70 and RPA32 (**Fig. 3b – e** & **Supplementary Figure 8)**. These data reinforce the model that most of the DNA binding properties of RPA that are promoted through the DBDs are driven by ssDNA interactions.

**Fig. 3.**
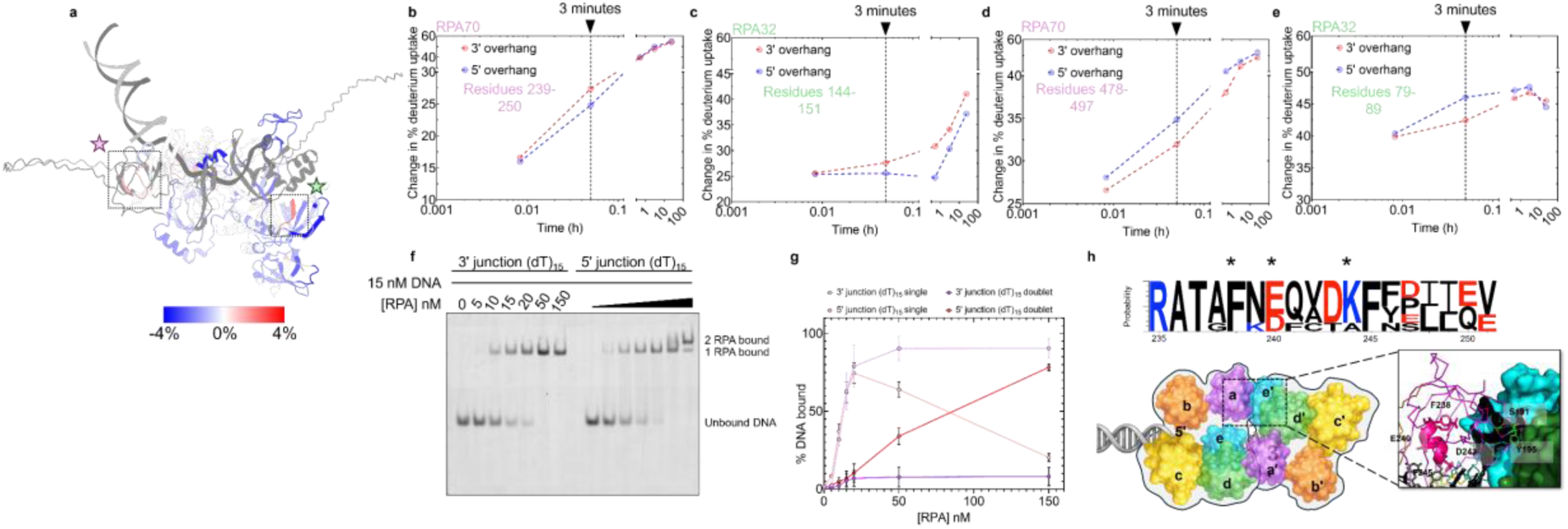
Conserved residues in RPA^DBD-A^ mediate junction-specific higher-order assembly with minimal global structural remodeling. **a,** AlphaFold3 model of an RPA-5ʹ junction (dT)_15_ substrate complex mapped with a percentage change in deuterium uptake (after 3 minutes of incubation) relative to the 3ʹ junction (dT)_15_ complex. Inset with magenta, and green stars represent regions with the highest uptake in RPA70, and RPA32 subunits, respectively. **b - e**, Scatter-plots representing the change in percentage deuterium uptake as a function of time for the RPA70 **(b and d)**, RPA32 **(c and e)** regions of interest (numerically highlighted within each graph). Dotted black line represents the three-minute time-point that was used to map changes on the AlphaFold3 structure. **f**, Electrophoretic mobility shift assay depicting RPA binding to 3ʹ and 5ʹ junction (dT)_15_ overhangs. **g**, Quantitation of the % DNA bound species for the single-bound and two-bound complexes as a function of RPA concentration between both junction conditions. Mean values connected, and the raw data from three independent repeats are depicted. **h**, Sequence conservation map depicting the degree of conservation within residues in the junction-proximal motif of RPA DBD-A. Taller letters indicate higher probability of conservation across multiple species. Asterisks (*) highlight residues that were chosen for characterization. Inset of the crystal structure of *Ustilago maydis* RPA bound to ssDNA (PDB: 4GNX), highlighting the implicated residues and the interaction between adjacent RPA molecules in the unit cell.

### Junction-specific higher-order assembly of RPA

The smFRET data and the preferential junction binding observations in the CTFM with respect to the 3ʹ junction agree and can be readily explained by the intrinsic stability of RPA at this position (**Fig. 2e**). In contrast, the behavior at the 5ʹ junction is more complex as RPA is stably bound in the CTFM assay, but the smFRET results suggest dynamic configurational rearrangements. Furthermore, the HDX-MS experiments show no robust structural changes specific to one junction over the other. The smFRET and HDX-MS investigations were carried out under experimental conditions where only one RPA is engaged on the DNA, however, the longer length of the DNA in the CTFM setup reflects a scenario where multiple RPA are loaded. We hypothesized that an RPA molecule at the junction might be influenced by trailing RPA molecules that solely engage the ssDNA. To test this hypothesis, we performed an electrophoretic mobility shift assay (EMSA) on DNA substrates containing a 5ʹ or 3ʹ junction along with a (dT)_15_ ssDNA overhang. Adding increasing amounts of RPA to the 3ʹ junction substrate produced a single shifted complex, and in agreement with a single RPA molecule stably bound at this position (**Fig. 3f**). In contrast, both one-RPA and two-RPA bound complexes are observed on the 5ʹ junction containing substrate (**Fig. 3f & g**). These data showcase a phenomenon at the 5ʹ junction, where configurational remodeling of the RPA bound at this junction paves way to the loading of additional RPA molecules on the trailing ssDNA (**Fig. 3h**). An AlphaFold3 model of two RPA molecules bound to DNA containing a 5ʹ junction-(dT)_15_ overhang shows a complex arrangement of the DBDs and PIDs (**Supplementary Figure 9**). DBD-A of the junction-bound RPA is positioned to interact with both RPA32 and RPA14 of the trailing RPA. In fact, the crystal structure of *Ustilago maydis* RPA shows interactions promoted by DBD-A and DBD-D & RPA14 from two neighboring DNA-bound RPA molecules in the unit cell (**Fig. 3h**). Based on these structures we propose that this RPA binding interplay collectively renders higher stability to the RPA molecules at the 5ʹ junction.

### Contributions from conserved, charged and aromatic residues in RPA^DBD-A^ to junction-specific engagement

In our HDX-MS analysis, residues 239-250 in DBD-A of RPA70 show small changes in deuterium uptake (**Fig. 3a, b)**. This region resides at a conserved inter-RPA interface in the *U. maydis* RPA crystal structure (PDB: 4GNX; **Fig. 3h**). We also note that residues 185-194, that harbor a motif for phosphorylation in RPA70, is also situated in the interaction interface between the two RPA molecules. Therefore, we probed the role of conserved residues in this region through mutagenesis and investigation of junction-binding properties. The role of three highly conserved hydrophobic/charged residues (RPA70^F238^, RPA70^E240^, and RPA70^D243^) were screened (**Fig. 4**). All three recombinant mutant RPA variants bound to a (dT)_15_ ssDNA with affinity similar to wild-type RPA (**Supplementary Fig. 10a**). Notably, RPA70^F238A^ has a modest binding defect, relative to the other variants as detected in the band-shift assays. Next, we tested binding to the 5ʹ junction (dT)_15_ substrate. We observed a marked reduction in binding affinity for the mutants compared to wild-type RPA, suggesting that the mutations impair junction-specific engagement (**Supplementary Fig. 10b**). Furthermore, we probed the effects of these residues on the facilitated exchange and initial engagement assays. In the facilitated exchange set up, we found that the mutants showed a marked reduction in their residence time ratios relative to the wild-type condition, suggesting distinct junction-proximal engagement dynamics (**Fig. 4a-d; Supplementary Fig. 11a-13a**).

**Fig. 4.**
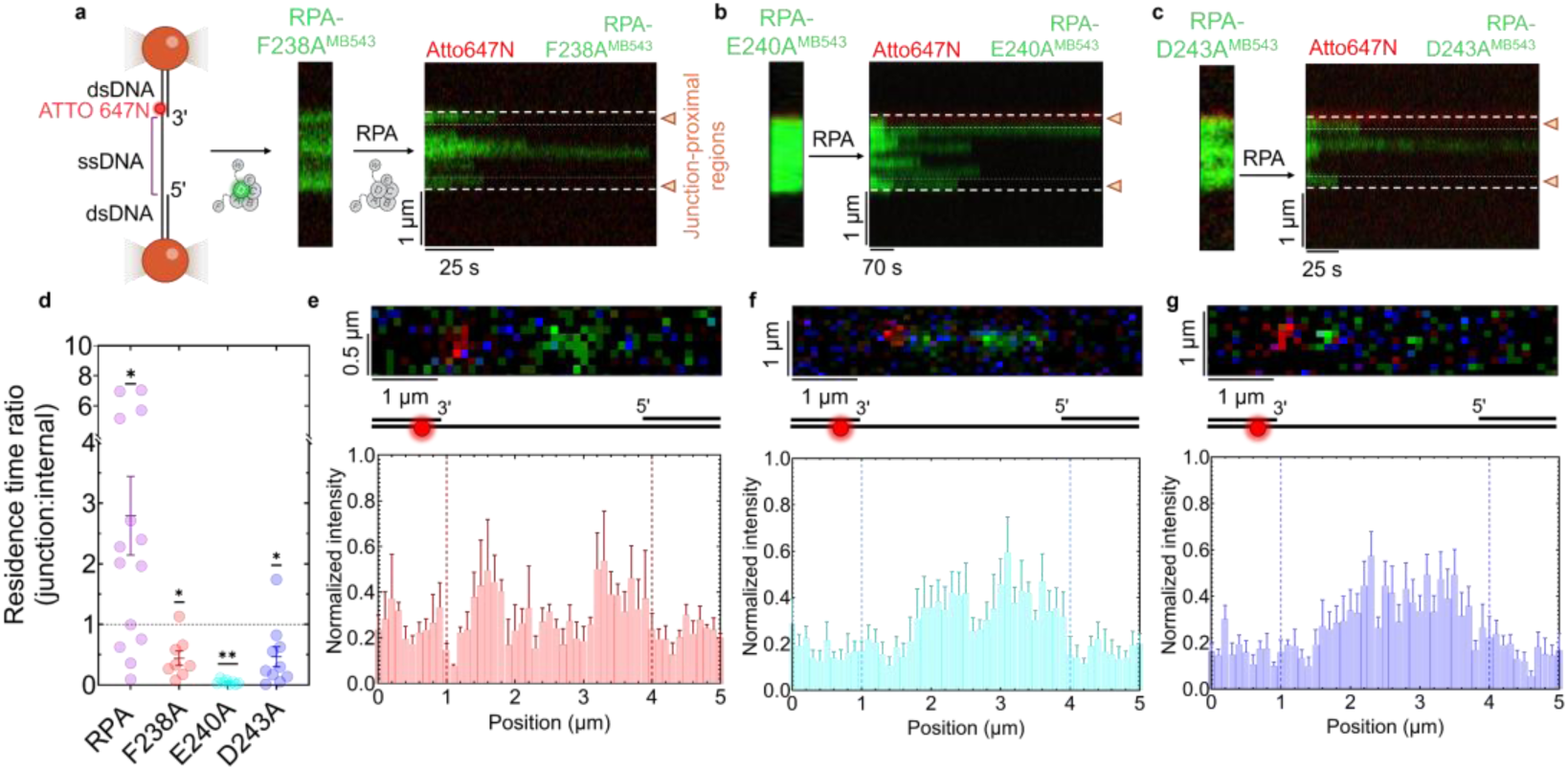
Regulation of junction-specific engagement dynamics by the conserved RPA DBD-A motif. **a - c**, Representative kymographs capturing facilitated exchange across the fluorescently labeled RPA-F238A **(a)**, E240A **(b)**, and the D243A **(c)** mutants on the gapped DNA substrate. **d**, Dot-plot indicating junction-proximal: internal RPA^MB543^ residence time ratios plotted relative to a baseline ratio of 1 (indicating equivalent residence times); mean ± SEM is depicted, each data point is a single junction: internal residence time ratio. The non-parametric one sample Wilcoxon test was applied to compare the median with a fixed median value of 1. (*) indicates p-value ≤ 0.033, and (**) ≤ 0.0021. Data collected from N= 8-14 binding events. **e - g**, Representative image scans of the first frame capturing detectable binding of the fluorescently labeled RPA-F238A **(e)**, E240A **(f)**, and D243A **(g)** mutants. Bar graphs of normalized intensities for this initial binding signal are plotted as a function of position along the gapped DNA substrate under each mutant image scan. The dotted lines represent the ss/dsDNA junction, and junction polarity is defined by the position of the Atto647N dye (shown in red). Mean ± SEM plotted for each condition.

Spatially mapping the initial binding events pointed to a modest reduction in junction-proximal initial binding for all the mutants (**Fig. 4e-g; Supplementary Fig. 11b-13b**). While the binding profile across multiple DNA tethers was comparable to the wild-type condition for RPA70^F238A^ in terms of junction-proximity (**Fig. 4e; Supplementary Fig. 11b**), it was evidently more promiscuous for the RPA70^E240A^ and RPA70^D243A^ mutants (**Figure 4f, g; Supplementary Fig. 12b, 13b**). These findings suggest that residues within the DBD-A motif identified are indeed contributing to the overall regulation of RPA’s junction-specific engagement dynamics. RPA70^F238^ could base-stack with the terminal exposed base at the junction, while RPA70^E240^ and RPA70^D243^ could directly hydrogen bond with individual bases at the junction (**Supplementary Figure 9**). Alternatively, these residues could promote a specific configuration of the RPA domains through inter- or intra-RPA protein interactions.

### RPA phosphorylation dampens junction-specific engagement dynamics

RPA phosphorylation is a key post-translational modification (PTM) observed across multiple cell-cycle stages and DNA metabolic events^8,33–37^. We have previously shown that ssDNA binding does not change the overall configurational spacing of the DBDs or the PIDs^21,22^, but PTMs such as phosphorylation release the PIDs to promote protein-protein interactions^23^. RPA phosphorylation is also known to reconfigure the global, domain-level arrangement of RPA’s OB-folds^22,23,38^. A key observation in our earlier studies of RPA phosphorylation was the binding of a higher density of phospho-RPA molecules^39,40^. Given the position of T191 (which is phosphorylated as part of a phosphocode during G2-M) in the two-RPA bound context (**Supplementary Fig. 14**), we hypothesized phosphorylation might also affect RPA’s junction-specific engagement dynamics.

To test this hypothesis, we used two RPA phospho-mimetic variants: RPA^2D^ and RPA^3D^. RPA^2D^ carries two phosphomimetic Asp substitutions at positions Ser23 and Ser29 in the N-terminus of RPA32. These are known basal and priming sites of phosphorylation that occurs in both the S, G2, and mitotic phases of the cell-cycle^8,29,33–35^. RPA^3D^ possesses one additional phosphomimetic Asp substitution at position Thr191 in the large RPA70 subunit that is specific to G2-M. We performed the facilitated exchange and initial binding assays and uncover reduced residence time ratio for both RPA^2D^ and RPA^3D^ during facilitated exchange (**Figure 5a-c; Supplementary Fig. 15a, 16a**). Interestingly, between the two phosphomimetic forms, RPA^3D^ displays relatively more rapid eviction, suggesting functional differences for these RPA phosphocodes (**Figure 5c; Supplementary Figure 16a**). Both phosphomimetics demonstrate a slight preference to initiate engagement proximal to the junctions. While RPA^2D^ displays a preference for the 3ʹ junction (**Fig. 5d; Supplementary Fig. 15b**), RPA^3D^ showed a marked preference for the 5ʹ junction on the gapped DNA substrate (**Fig. 5e; Supplementary Fig. 16b**). These data suggest that RPA’s junction-proximal engagement dynamics appear to be regulated by their phosphorylation status and tuned for junction selectivity.

**Fig. 5.**
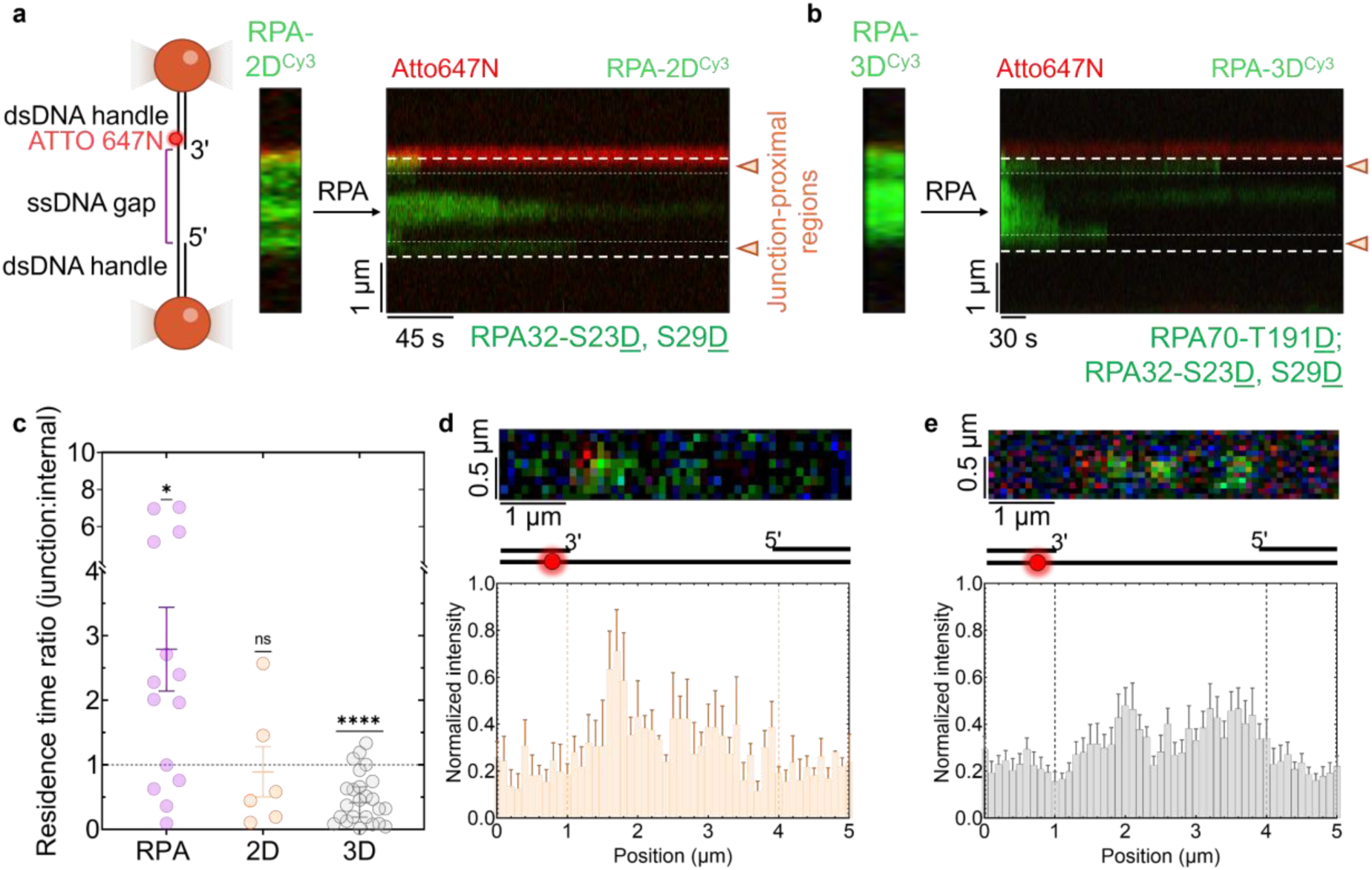
RPA phosphorylation alters junction-specific engagement dynamics. **a, b**, Representative kymographs highlighting the loss of RPA-2D^Cy3^ **(a)** and RPA-3D^Cy5^ **(b)** (0.25-0.5 nM) when challenged with excess (2.5-5 nM) unlabeled RPA in channel 5. **c**, Dot-plot indicating junction-proximal: internal RPA^Cy3/Cy5^ residence time ratios plotted relative to a baseline ratio of 1 (indicating equivalent residence times); mean ± SEM is depicted, each data point is a single junction: internal residence time ratio. The non-parametric one sample Wilcoxon test was applied to compare the median with a fixed median value of 1. (**) indicates p-value of ≤ 0.0021. Data collected from N= 6-24 binding events. **d, e**, Representative image scans of the first frame capturing detectable binding of the fluorescently labeled RPA-2D and RPA-3D variants. Bar graphs of normalized intensities for this initial binding signal are plotted as a function of position along the gapped DNA substrate for RPA-2D **(d)** and RPA-3D **(e)**. The dotted lines represent the ss/dsDNA junction, and junction polarity is defined by the position of the Atto647N dye (shown in red). Mean ± SEM plotted for each condition.

### RPA regulates XPA’s junction-specific recruitment and engagement dynamics

We have previously shown that the recruitment of the *Saccharomyces cerevisiae* homologous recombination Rad51-Rad52-Rad55/57 complex on a gapped DNA substrate is spatially coordinated at the 5ʹ junction by Rpa^16^. Here, we test and extend the idea to human RPA and an RPA-interacting protein XPA – a nucleotide excision repair (NER) pathway protein. In the CTFM setup, we first tested the binding of XPA^Cy5^ to the gapped DNA substrate (**Fig, 6a; Supplementary Fig. 17b**). We found that XPA^Cy5^ showed frequent, yet isolated and short-lived binding events promiscuously along the 5005-nt ssDNA region (**Fig. 6a, b**). Remarkably, when we coated the gap with RPA^MB543^ and then introduced XPA^Cy5^, we saw that XPA^Cy5^ preferentially localized to the 5ʹ junction-proximal region and demonstrated a ∼30-fold increase in the average binding lifetime (**Fig. 6c-f; Supplementary Fig. 17a, c**). Next, we tested the effect of a junction-specific RPA mutant (F238A) on XPA recruitment. Consistent with our hypothesis, XPA^Cy5^ failed to localize at the junction when the gap was coated with RPA-F238A^MB543^ (**Fig. 6e**). A recovery in XPA^Cy5^ binding lifetime was observed, suggesting that RPA-F238A was specifically impaired at its junction-specific recruitment function, while permitting interactions with XPA in the non-junction, internal ssDNA regions (**Fig. 6f**). Collectively, these findings point at the indispensable role of junction-proximal RPA and the junction-specific motif, in this exemplar case at the 5ʹ junction, in governing where XPA is recruited, and how stably it remains at that site.

**Fig. 6.**
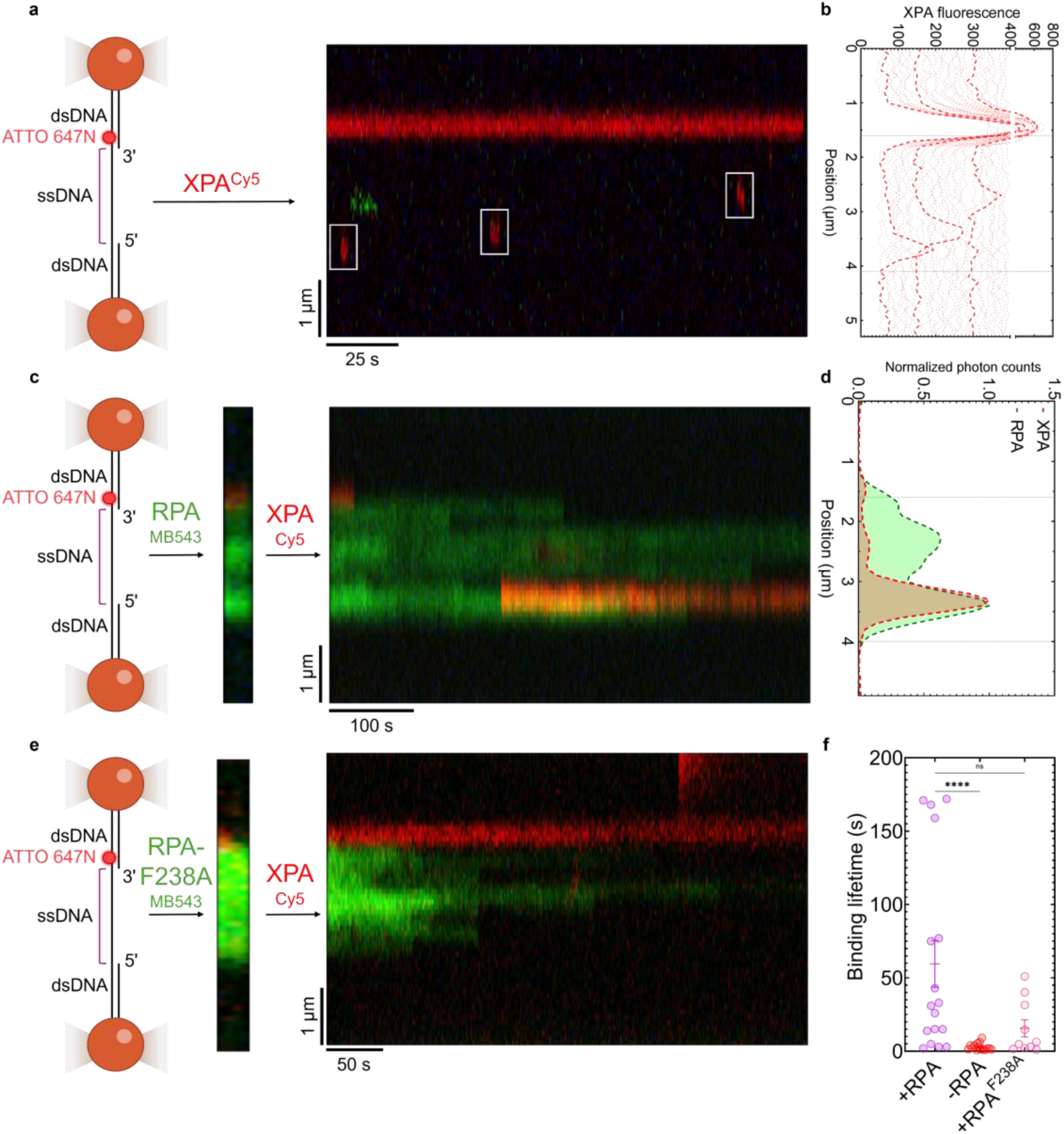
Junction-proximal RPA directly regulates XPA recruitment and binding lifetimes on gapped DNA substrates. **a**, Representative kymograph highlighting XPA^Cy5^ binding to a gapped DNA substrate. **b**, Signal quantification of XPA^Cy5^ as a function of position on the gapped DNA substrate. Lines indicate average signal binned for every ∼35 line scans. **c**, Representative kymograph depicting XPA^Cy5^ binding to an RPA^MB543^ coated gapped DNA substrate. **d**, Normalized XPA^Cy5^ and RPA^MB543^ photon counts for the kymograph in (**c**) as a function of position along the gapped DNA substrate. Dotted lines demarcate the ss/dsDNA junction regions., Representative kymograph depicting XPA^Cy5^ binding to an RPA^MB543^ coated gapped DNA substrate. **e**, Representative kymograph depicting XPA^Cy5^ binding to an RPA-F238A^MB543^ coated gapped DNA substrate. **f**, Dot-plot indicating the binding lifetime (s) for XPA^Cy5^ on a gapped DNA, RPA-coated gapped DNA, and RPA-F238A coated gapped DNA substrate. Mean ± SEM plotted for each condition, and the one-way ANOVA test with Dunn’s correction was applied to compare the conditions. (**) indicates p-value of ≤ 0.0021. N= 10-17 binding events were compared across all conditions.

## Discussion

Binding site specificity for DNA processing enzymes is canonically informed by sequence or structure recognition and is prominent in the context of double-stranded DNA^41^. However, replication, recombination, and in most DNA repair scenarios, ssDNA serves as the template and is completely sequestered by RPA and buried within its high-affinity ssDNA binding domains ^42,43^. However incoming DNA metabolic proteins must engage the ssDNA, both in terms of spatial and temporal specificity, for successful progression of DNA replication or repair. How and where these interactions occur on the RPA-coated DNA, and the underlying mechanisms of DNA handed off remains unknown. Here, we uncover an important mechanistic feature for RPA showcasing preferential initial engagement and higher stability at junction-proximal regions. These DNA binding properties are also different from the well-understood behavior of RPA on ssDNA. The mechanisms underlying these junction-specific features also appear to be unique at the 5ʹ versus 3ʹ junctions and point to the evolution of functional specificity.

Our data show that RPA preferentially engages the two junctions, but using independent mechanisms. At a 3ʹ junction, stable binding of RPA is likely enacted by its DNA melting property. The directional movement of RPA has been shown to unwind short hairpin structures but is strictly coupled to a 5′->3′ directionality on the ssDNA overhang. Therefore, at the 3′ junction, we envision RPA invading a short dsDNA region (**Fig 7d**). In contrast, the stability at the 5′ junction must invoke a mechanism that is independent of dsDNA melting. Based on available structures, and our observation of higher-order RPA complexes formed on the 5′ junction containing substrates, we propose that stability and specificity are driven through inter-RPA interactions between two RPA molecules at this junction (**Fig 7a)**.

**Fig. 7.**
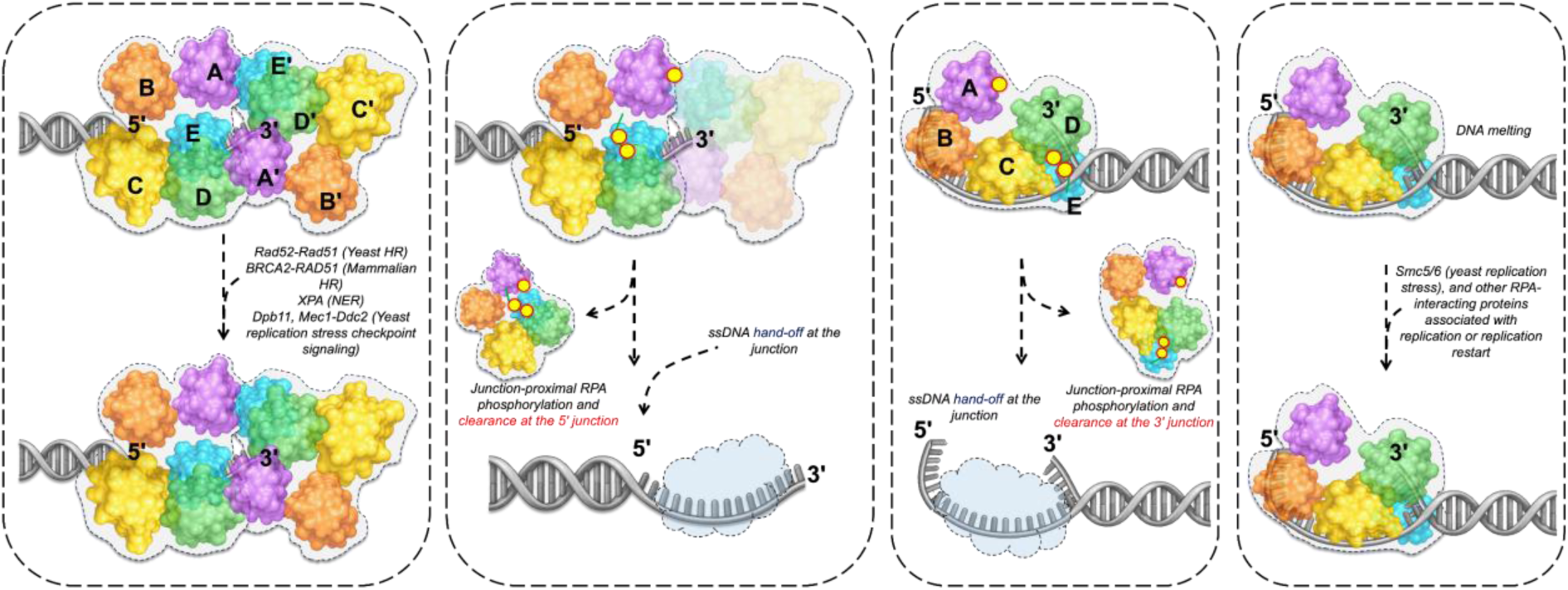
Proposed models for junction-proximal RPA-driven regulation of diverse DNA metabolic events. Transient ssDNA intermediates with a defined ss/dsDNA junction arise during most DNA metabolic events such as replication, repair and recombination. The model depicts assembly and cooperation between two RPA molecules at the 5ʹ junction. At the 3ʹ junction, the model posits that stable binding of RPA arises from the transient melting of the dsDNA. The unique configuration of the RPA molecule at the junctions is proposed to drive recruitment of specific RPA-interaction proteins to the junctions. Phosphorylation of RPA molecules bound to the junction results in the rapid clearance of RPA and thus enabling efficient ssDNA hand-off to incoming DNA binding proteins.

In the cell, RPA-occluded ssDNA that are adjacent to 5ʹ versus 3ʹ junctions would be encountered during different scenarios of DNA metabolism. In the context of homologous recombination (HR), end-resection results in the formation of a 3ʹ overhang substrate with a 5ʹ junction. HR and microhomology mediated end-joining (MMEJ) are two examples where incoming proteins need to function on 5ʹ junction containing DNA carrying 3ʹ ssDNA that are occluded by RPA. Recent work on assembly and recruitment of mediator-recombinase complexes onto RPA-coated DNA during HR show initial recruitment at the junction in both mammalian and yeast HR systems^44,45^. This recruitment is strictly dependent on physical interaction between RPA and the mediator proteins. Similar recruitment between RPA and the Polymerase θ (Polθ) protein has been captured during MMEJ^18,19^.

Long stretches of 3′ ssDNA overhangs are thought to be coated by RPA during HR in contrast to shorter ssDNA overhangs found during MMEJ. Therefore, the proposed models for junction binding are likely independent of ssDNA length. At the 5′ junction, irrespective of the length of the ssDNA overhang, we propose that configurational rearrangements of the highly dynamic DBDs-A & B give rise to open space on the ssDNA for binding of additional RPA molecules. Stable binding of RPA is likely driven by the less-dynamic Tri-C complex composed of DBDs-C & D with OB-E serving as the anchor holding together the constitutive trimer^46^. Now, whether the configurations of the trailing RPA mirror those of the junction-bound RPA remain to be established. More importantly, how the protein-interaction domains are accessible within the junction-bound versus training RPA might be a key determinant of recruitment of an incoming protein as shown by XPA interactions in this study.

RPA phosphorylation is widely observed across different cell cycle stages and DNA metabolic contexts. RPA is phosphorylated at the N-terminus of RPA32 and at various positions on RPA70. Defined combinatorial phosphocodes have been identified specific to both G2-M and in mitosis. HR is suppressed in mitosis through phosphorylation of RPA. The same modification could also in principle support MMEJ in mitosis. Given the junction-specific localization of the HR mediator-recombinase complex and the MMEJ-specific Polθ to 5ʹ junction, we tested where phosphomimetic versions of RPA would show altered junction-binding properties. We previously showed that RPA carrying two phosphomimetic substitutions at S23 and S29 positions in RPA (RPA^2D^), along with an additional substitution at T191 in RPA70 (RPA^3D^) have no appreciable differences in ssDNA binding^39^. Nonetheless, both RPA^2D^ and RPA^3D^ showed vast configurational rearrangements within the OB-folds, especially the protein-interaction domains^39^. In terms of junction recognition properties, RPA^3D^ had a slight preference to initiate engagement proximal to the 5ʹ junction, while RPA^2D^ displayed a similar preference for the 3ʹ junction. Interestingly, both RPA^2D^ and RPA^3D^ displayed an inability to remain stably bound at the junctions when challenged with excess RPA. RPA^2D^ was relatively more stable than RPA^3D^ in terms of exchange, although it wasn’t significantly more stable relative to the internally bound RPA molecules. Both these observations suggest that RPA phosphorylation, in a context-specific manner, can fine tune its junction-specific engagement dynamics, which we posit can affect how RPA recruits specific downstream partners. These findings also provide a mechanism by which RPA can be removed from the junctions post recruitment of partner proteins to hand-over the DNA (**Fig. 7b, 7c**). DNA repair and cell-cycle associated kinases such as ATR, AurkB, and CDK1 have all been shown to phosphorylate RPA and in turn are regulated by RPA through a feedback mechanism^38,47,48^. In yeast, Dpb11 binding and the subsequent recruitment of the Mec1-Ddc2 complex to the 5ʹ junction are directed by RPA^14^. The mechanisms regulating RPA-kinase interactions and their direct functional roles remain to be uncovered.

The significance of junction binding is highlighted by the RPA mutations in DBD-A. All three RPA mutants display impaired junction stability. Our observations of the mutants specific to junction-engagement correlate with phenotypes observed *in vivo*. For example, an RPA-F238A substitution is lethal in *Saccharomyces cerevisiae*^49^. A homologous substitution in human RPA acts as a separation-of-function mutant, resulting in faithful DNA replication, but impaired DNA repair, and consequent cell-cycle arrest^50^. Our work now provides a mechanistic basis for these phenotypic endpoints. The behavior of RPA at the 3ʹ and 5ʹ junction points at potential implications for junction-specific DNA metabolic events. For examples, recent work implicate RPA in the junction-specific recruitment of Polθ and the regulation of theta-mediated end joining at the 5ʹ junction during MMEJ^18,19^. Our observations of a two RPA bound 5ʹ junction substrate carrying a (dT)_15_ overhang aligns with the new structural work with Polθ, where destabilization of DBDs C & D of RPA by Polθ likely tilts ssDNA binding to Polθ’s favor, driving the hand-off.

In biology, 3ʹ junctions would be encountered in processes such as DNA replication, particularly on the lagging strand Okazaki fragments. Almost all DNA repair processes that depend on DNA replication to complete strand synthesis would also encounter a 3ʹ junction. In the context of replication stress, recent studies show recruitment of the Smc5/6 complex to the 3ʹ junction in an RPA and ATP-dependent manner^17^. Another key example for 3′ junction binding and melting activities of RPA arises from recent work on the Pif1 helicase where no unwinding was observed in the absence of RPA. But RPA engagement at the 3′ junction and subsequent transient melting enables a lagging Pif1 molecule to propel the RPA further into the duplex leading to unwinding. This functionally fits with Pif1’s role in unwinding 3′ junctions arising at stalled replication forks. Our observations with RPA and XPA further situate RPA’s role within the context of NER.

XPA is understood to work in tandem with RPA during damage recognition in NER, upstream of recruiting the larger NER complex to the site^51^. However, the precise role of RPA in regulating XPA’s recruitment and stimulating its activity was unknown. Based on our findings that RPA significantly regulates the spatial recruitment and average binding lifetimes of XPA at the 5ʹ junction, we point at the role of RPA in stably situating XPA at the correct position along DNA to coordinate the timely and effective recruitment of downstream NER-specific protein complexes, like the XPF-ERRC1 complex to the repair site^52–54^.

Overall, our work informs our understanding of RPA on two inter-connected aspects. First, it reinforces the conceptual framework around recognizing RPA’s engagement of ssDNA substrates in a context-dependent manner. RPA binding to ssDNA is not a static, protective measure alone, but rather is an appreciably dynamic endeavor that can regulate a range of downstream events. Second, and relatedly, RPA’s distinct ability to engage a overhang substrate at the junction-proximal region enables us to identify the basis of junction-specificity for a variety of incoming DNA metabolic proteins.

## ACKNOWLEDGEMENTS

This work was supported by funding from the National Institutes of Health - National Institute of General Medical Science R35-GM149320 (E.A.); the National Cancer Institute R01-CA305375 (E.A.), the Office of the Director S10-OD030343 (E.A.). A.M. acknowledges tuition support from the Philip and Lillian Katzman Scholarship, and the Doisy Scholarship from the Department of Biochemistry and Molecular Biology, St. Louis University School of Medicine.

## AUTHOR CONTRIBUTIONS

Conceptualization: EA, RC, & AM

Methodology: AM, RC, EH, BBB, & EA

Investigation: AM, RC, EH, AA, MJ, IK, TS, & MTL

Visualization: AM, EH, & EA

Funding acquisition: EA, & BBB

Project administration: EA

Supervision: EA & BBB

Writing – original draft: AM & EA

Writing – review & editing: all authors

## COMPETING INTERESTS

The authors declare no competing interests.

## DATA AVAILABILITY

The hydrogen-deuterium exchange mass spectrometry proteomics data have been deposited to the ProteomeXchange Consortium via the PRIDE partner repository with the dataset identifier PXD087967.

## CODE AVAILABILITY

The Python scripts used to process kymographs, and scripts used for analyzing smFRET data are available at the following GitHub repositories - https://github.com/AyushMistry25, and https://github.com/ChaddaRah/Single-Molecule-Trajectory-Analyzer_RahulChadda.

## METHODS

### Purification and labeling of RPA, non-canonical amino acid-carrying RPA, and phosphomimetic RPA variants

Unlabeled and fluorescently labeled human RPA, RPA phosphomimetics (hRPA70-32^S23D-S29D^-14 (2D), and hRPA70^T191D^-32^S23D, S29D^-14 (3D)) and *Saccharomyces cerevisiae* Rpa were purified as described^21,26,55,56,39^. For the optical trap experiments, RPA carrying MB543 on DNA-binding domain D was used. Primary amine labeling for RPA-2D and RPA-3D was performed in the following manner. Following overnight dialysis in the labeling buffer (30 mM Tris-HCl [pH 7.0], 200 mM KCl, 0.25 mM EDTA [pH 8.0], 1X protease inhibitor cocktail (PIC), and 10% v/v glycerol), the protein was incubated with two-fold excess NHS-Cy3 for 90 minutes at 4°C with gentle rocking. The reaction was then passed through a Biogel P-4 column (Bio-Rad Inc.) equilibrated with storage buffer (30 mM HEPES [pH 7.8], 200 mM KCl, 0.01% Tween-20, 1X PIC, 0.2 mM EDTA [pH 8.0], and 10% v/v glycerol), and the pooled fractions were concentrated, pooled, flash-frozen using liquid nitrogen, and stored at -80°C. The final protein concentration and labeling efficiency were calculated spectroscopically using ɛ_280_ = 87, 410 M^-1^ cm^-1^ for the protein, and ɛ_550_ = 150, 000 M^-1^ cm^-1^ for NHS-Cy3, as well as a correction factor at 280 nm of 0.073.

### Correlative optical-trap experiments and analysis

Prior to use, all five channels of the microfluidics chip were passivated according to Lumicks protocol with minor modifications. Briefly, 0.1% BSA, 0.5% BSA, and 0.5% Pluronic F-127 solutions were sequentially flowed at 2 bar for 2 minutes, and 0.5 bar for 12 minutes, each. The solutions were allowed to incubate in the channels with no flow for 15 minutes. Finally, reaction buffer (30 mM HEPES [pH 7.8], 100 mM KCl, 5 mM MgCl_2_, and 0.2 mg/mL BSA) was flowed at 2 bar for 2 minutes, and 0.5 bar for 12 minutes, followed by incubation for 15 minutes with no flow to equilibrate the microfluidics chip. Biotinylated ds/ss DNA hybrid precursor (17.852 kbp, Cat #00027 Lumicks Inc.) and 4.35 µm Streptavidin-coated polystyrene beads from Lumicks were diluted in 1X running buffer (137 mM NaCl, 2.7 mM KCl, 1.12 mM phosphates, 5 mM sodium azide, and 0.5 mM EDTA, pH 7.4) according to manufacturer’s instructions. The DNA hybrid precursor was captured between two beads at 0.24-0.28 pN/nm trap stiffness, and ss DNA gap was generated by pulling the DNA in a low-salt buffer (0.1× running buffer). Formation of ssDNA gap was verified by comparison to the worm-like chain model in the instrument-associated Bluelake software.

Prior to visualization, the fluorescent and unlabeled proteins were diluted in reaction buffer to allow addition into the final reaction volume. For the facilitated exchange experiments, the ss/dsDNA hybrid was stretched between two beads with a force of 10 pN at a flow pressure of 0.05 bar. The DNA was moved to a microfluidics flow channel containing 0.25-0.5 nM MB543-labeled RPA variant(s) in the imaging buffer (reaction buffer supplemented with 2–3 mM Trolox, 0.165 U/µL glucose oxidase, 2.17 U/ µL catalase and 0.8% (w/v) D-glucose monohydrate). After confirming the complete coating of the ssDNA gap with fluorescent RPA, the DNA was then moved to another microfluidics channel containing 2.5-5 nM unlabeled RPA. Confocal microscopy was carried out using green (561 nm) and red (638 nm) lasers, with a pixel size of 100 nm and a dwell-time of 2 ms per pixel. Stacks of frames (kymograph) recorded by continuous confocal scanning along the DNA axis over time were visualized using the Lumicks-developed Lakeview Pro software. For the initial engagement experiments, the same arrangement was used, but continuous image scans were recorded, starting from the low-salt channel (Channel 3) following confirmation of hybrid DNA substrate, through entry into the first imaging channel (Channel 4).

All kymographs (and image scans, when necessary) were exported as.h5 files and processed using specific Python scripts on a Jupyter notebook. Kymographs were cropped to the DNA tether and calibrated from pixel index to genomic position using the known tether length (17, 853 bp). A percentile-based contrast scaling method was used to adjust the display. Background was estimated as the mean photon count within a manually selected region of the kymograph containing no RPA signal and was subtracted from every pixel of the image. A region of interest encompassing a binding event was selected, and the background-corrected photon counts were summed across all pixels in the region of each scan line, deriving total photon counts as a function of time. Binding lifetimes were derived by measuring the time taken from an increased photon count signal to return back to baseline for each binding event. For visual interpretation of noisy traces, a Savitzky-Golay filter was applied, and smoothed traces were derived for visualization. Raw and smoothed traces were exported as CSV files. The minimum quantifiable distance along the line-scan (∼0.3 µ/ 300 nm, based on the microscope’s diffraction limit of ∼200-250 nm) was taken from both junctions towards the ssDNA region of the hybrid substrate and designated as the junction-proximal region.

Positional binding along the DNA tether upon initial engagement was quantitated by analyzing the image scan frame that first captured a binding event. The frame was converted to arrays of photon counts indexed by pixel position along the tether. Background was similarly estimated as the mean photon count within a manually selected signal-free region of the scan and subtracted from every pixel. A region of interest, spanning the end-to-end length of the DNA tether was demarcated and photon counts were averaged across the selected frames for each pixel row. Pixel indices were subsequently converted using the pixel size of the measurement (0.1 μ), and profiles were exported as CSV files.

### TIRF / smFRET

smFRET-based analysis of RPA binding on immobilized (dT)_15_ overhang DNA substrates were carried out by injecting the fluorescently labeled RPA i.e. 100-150 pM of each RPA-D^Cy3^ and/or RPA-A^Cy5^ into an optical cell built over an inverted, objective based total-internal-reflection fluorescence microscope (TIR-FM; IX71 Olympus). TIR excitation was achieved through an oil-immersion objective (100X Olympus UplanApo Numerical Aperture 1.5). The sample was excited with a 532 nm laser, and the emitted fluorescence was split into two channels using Optosplit II (Cairn-Research, UK); Cy3 and Cy5 emissions were collected one half of the chip of the same electron-multiplying charge-coupled device (EM-CCD) camera (iXon Ultra DU-897U-CS0). Images were acquired with 150 ms frame time. Donor and Acceptor images were subpixel registered in Fiji using an image of multicolor beads as fiducial markers. Next, background corrected intensity traces were extracted from immobilized single-molecule spots using Matlab scripts. FRET values were calculated for each fluorescent spot, namely ratio between the acceptor intensity and the sum of the intensities of the donor and acceptor, corrected for crosstalk and cross-excitation. The single molecule FRET v time traces were analyzed in vbFRET to ascribe FRET states and dwell times.

Glass coverslips for single-molecule imaging were cleaned and coated as follows: Briefly No. 1.5 thickness Gold Seal coverslips (22 x 60 x 0.17) were sonicated in deionized water, 2 % Micro-90, 200-proof ethanol, and finally in 1 N KOH all at 60 °C for 15 min. each interspersed with exhaustive washes with deionized water. After drying under a stream of filtered N_2_, the coverslips were first coated with Vectabond (1 % v/v Vectabond in 1:18 methanol/acetic acid mixture) followed by washing with deionized water. Next, dried coverslips were incubated overnight with a freshly prepared 18:1 mixture of mPEG-SVA and Biotin-PEG-SVA in 0.1 M NaHCO_3_. Nonspecific binding, which was measured by flowing in non-biotinylated Cy3- or Cy5-labeled DNA or by flowing in biotinylated fluorescent DNA substrate in absence of neutravidin, was < 1 %. All measurements were performed at room temperature in buffer (50 mM Tris-acetate pH 7.5, 50 mM KCl, 5 mM MgCl_2_, 10 % v/v glycerol, 1 mM DTT, and 0.2–0.5 mg/mL BSA) supplemented with oxygen scavenging solution i.e. 0.1 mg/mL glucose oxidase, 0.2 mg/mL catalase, and 0.4 % (wt/wt) β-D-glucose (or 0.1 mM PCD / 10 mM PCA), and 2.5 mM trolox to reduce blinking. To a neutravidin (0.2 mg/mL) coated slide, biotin tagged (dT)_15_ overhang DNA substrates were added at 100–150 pM concentration to arrive at optimal single-molecule number-density of ∼ 1.2 spot per 3.5 mm^2^ area on the EM-CCD chip.

### Hydrogen-deuterium exchange mass spectrometry

HDX was performed with analysis of peptides via liquid chromatographs mass spectrometry (LC-MS) for RPA, RPA in complex with a 5’ (dT) substrate, and RPA in complex with a 3’ (dT) substrate, in triplicate. Initial protein concentration for all three conditions was 5.1 mg/mL. Exchange buffer was prepared by freeze-drying protein buffer without glycerol (30 mM HEPES, 200 mM KCl) and resuspending in 100% D_2_O (Thermo-Fisher). Proteins were diluted to a final concentration of 75% D_2_O at room temperature. After 30 seconds, 3 minutes, 30 minutes, 3 hours, and 24 hours, proteins were diluted fivefold into quench buffer (1 mg/mL porcine pepsin, 1% formic acid, 1.15M urea). Proteins were allowed to digest in quench buffer for 2 minutes on ice and were immediately flash-frozen in liquid nitrogen. Undeuterated samples were prepared in a similar manner, using undeuterated glycerol-free protein buffer.

LC-MS was performed with a 1290 UPLC series chromatography stack (Agilent Technologies) directly coupled to a 6538 Q-TOF mass spectrometer (Agilent Technologies) using a modified protocol that has been previously described^57^. To separate peptides, a reverse phase C18 column was used (Phenomenex Onyx Monolithic C18 column, 50 mm x 2 mm) at 1°C, with a flow rate of 500 µL/min. The following gradient was used: 0.0-0.7 min, 5% B; 0.7-4.5 min, 5-40% B; 4.5-5.5 min, 40-95%; 5.5-5.8 min, 95% B; 5.8-5.9 min, 5% B; 5.9-7.5min, 5% B. Solvent A was comprised of 0.1% formic acid in water (Thermo-Fisher) and solvent B contained 0.1 % formic acid in acetonitrile (Thermo-Fisher). Data acquisition was performed at 2.1 Hz over a scan range of 100-1700 m/z in positive mode. The electrospray settings were as follows: nebulizer 3.8 bar, drying temperature 350°C, drying gas 8.0 L/min., and capillary voltage 3.5 kV.

Peptides were identified prior to performing HDX by running a non-deuterated RPA sample seven times in auto MS/MS mode using an exclusion list. Data was collected with a 4 m/z isolation window at 1 Hz, with a precursor ion selection set to a maximum of five ions per cycle, sorted by abundance. Data was converted to mzML format via MS Convert ProteoWizard package version 3.023199 64 bit) using the vendor specific peak picking filter. Peptide identification was done via Peptide Shaker (CompOmics, version 3.0.8), selecting unspecific cleavage sites, precursor charge of 1-6, precursor tolerance of 50 ppm, fragment tolerance of 30 ppm, and oxidization of methionine as a variable modification using the Comet, Sage, and X!Tandem search algorithms. Identified retention times and protein sequences were used and further validated in subsequent HDX analysis. Deuterium uptake was determined by monitoring shifts of protein isotopic distributions in HDExaminer (Trajan Automation, version 2.5.1), using identified peptides and measured deuterated peptides. All deuterated and non-deuterated peptides were manually confirmed and deemed confident at a confidence value of 0.60 or higher. Using these parameters, 47 confident peptides were identified in RPA 70, 32 in RPA 32, and 21 in RPA 14, yielding 60%, 82%, and 99% sequence coverage overall (Supplementary figures ##). Percent deuterium incorporated (%D) was calculated using theoretical values provided from the HDExaminer Software. HDX uptake plots were exported from HDExaminer and have been provided as raw data.

The inter-junction percentage change in deuterium uptake was calculated for the 3-minute time-point by subtracting the 3ʹ junction value from the 5ʹ junction condition value for each peptide. The peptides showing the largest increase in percentage deuterium uptake were identified for each RPA subunit and were highlighted on the AlphaFold3 model of RPA and a 5ʹ junction-containing (dT)_15_ overhang substrate. In addition, the peptide covering 239-250 in RPA70, a region of interest, is shown. This peptide showed the second-largest increase in that subunit and was included on the basis of its functional relevance as well as its magnitude of change. Data are available via ProteomeXchange with identifier PXD087967.

### Measurement of ssDNA binding activity using EMSA

The assay was performed as descrribed^40^ with slight modifications. Briefly, 15 nM fluorescently labeled ssDNA (dT)_15_, or overhang (dT)_15_ substrates with either junction polarities were incubated with indicated amounts of RPA at room temperature in EMSA buffer (20 mM Tris-HCl [pH 7.8], 150 mM KCl, 5 mM MgCl_2_, 1 mM DTT, and 0.1 mg/mL BSA) for 15 minutes. The reaction mixture (30 μL) was mixed with 3 uL 70% v/v glycerol, mixed, and loaded onto a 5% non-denaturing TAE gel, and resolved using 0.5X TAE buffer at 4°C in the dark. The gels were pre-run for 30 minutes prior to sample loading, and the samples were resolved at 90 V over 100 minutes, at a constant voltage. The gels were scanned using an iBright 1500 imager (Thermo Fischer Inc.) and the fluorescence associated with ssDNA (unbound fraction) or the ssDNA + RPA (bound fraction) was background subtracted and quantified using the iBright Analysis Software (v. 5.3.0). The mean ± SEM from three independent experiments were plotted for analysis. K_D_ was estimated using the non-linear specific binding fit with Hill slope model (Eq. 1):

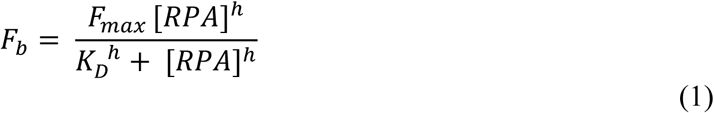

*F**_b_*** is the fraction of bound ssDNA determined from fluorescence intensities of two bands, namely:

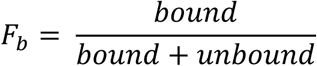

*F_max_* is the fraction of bound ssDNA at saturating protein concentration, *K_D_* is the apparent dissociation constant, *[RPA]* is concentration of RPA in each well, and *h* is the Hill coefficient.

### Structural Models

A model of RPA bound to the 5ʹ junction-containing (dT)_15_ overhang substrate was generated using the amino acid sequences for all three RPA subunits, and a defined oligonucleotide sequence to generate the partially annealed overhang using the web-service AlphaFold server powered by AlphaFold3^58^. Molecular graphics were modified using UCSF ChimeraX (v. 1.10), and the percentage change in deuterium uptake was modeled by converting the percentage change values to an attribute file format (.defattr extension), and labeling regions with their respective attribute values.

## Notes

### Competing Interest Statement

The authors have declared no competing interest.

### Summary of Updates

Revised Figure Legends Figure 7 and the associated Figure Legend were updated to better explain the model

