## Supplementary Information for "Replication Protein A recognizes DNA junctions to spatially organize genome maintenance"

**Supplementary Figures 1 – 17**

**Supplementary Tables 1 - 2**

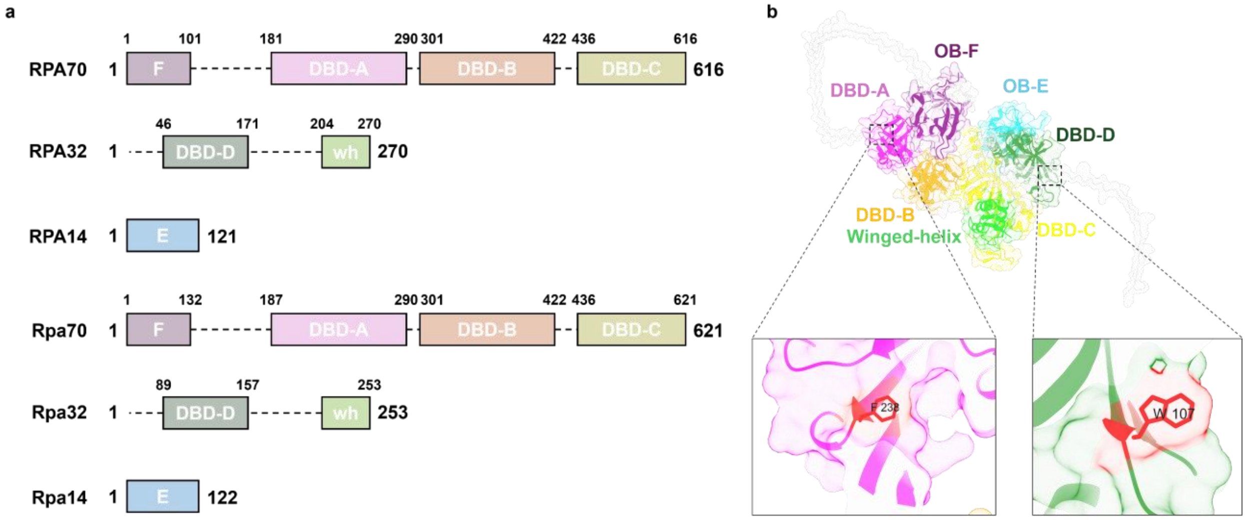


**Supplementary Figure. 1 | Domain organization of RPA, and the sites for ncAA-based fluorophore incorporation.**

**a,** Schematic representation of the 6 OB-folds within the human and yeast (denoted in lower case) RPA heterotrimer. The ordered domains are depicted in solid boxes, and the disordered linkers are shown as dotted lines. **b,** An AlphaFold3 model of human RPA, with the domains colored and annotated according to their corresponding residues. Inset within DBD-A and DBD-D highlight the residues that were fluorescently labeled via the ncAA-based labeling strategy for the optical trap and smFRET experiments. The position of the fluorophore had no effect on the junction preference or DNA binding properties of RPA.

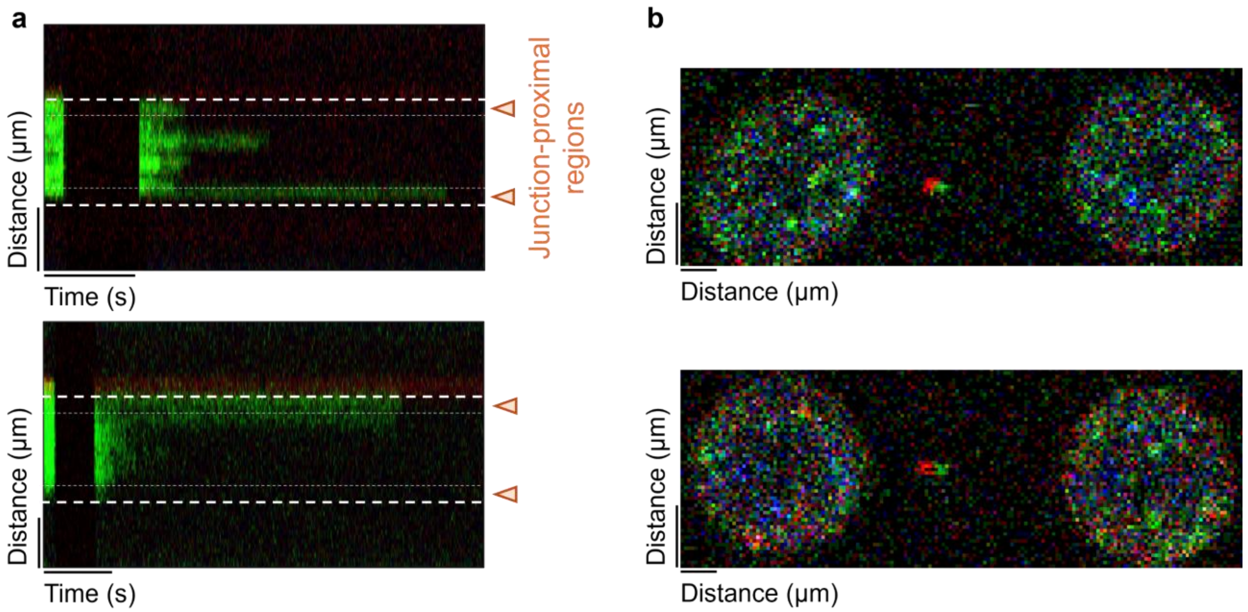


**Supplementary Figure. 2 | Facilitated exchange and initial engagement with hRPA.**

**a**, Representative kymographs depicting facilitated exchange with fluorescently labeled hRPA. **b**, Representative image scans depicting hRPA signal (in green) as it initiates binding. The red label represents an ATTO647N dye. The white dashed and dotted lines denote the area used to define and quantify ss/dsDNA junction & proximal regions in the kymograph.

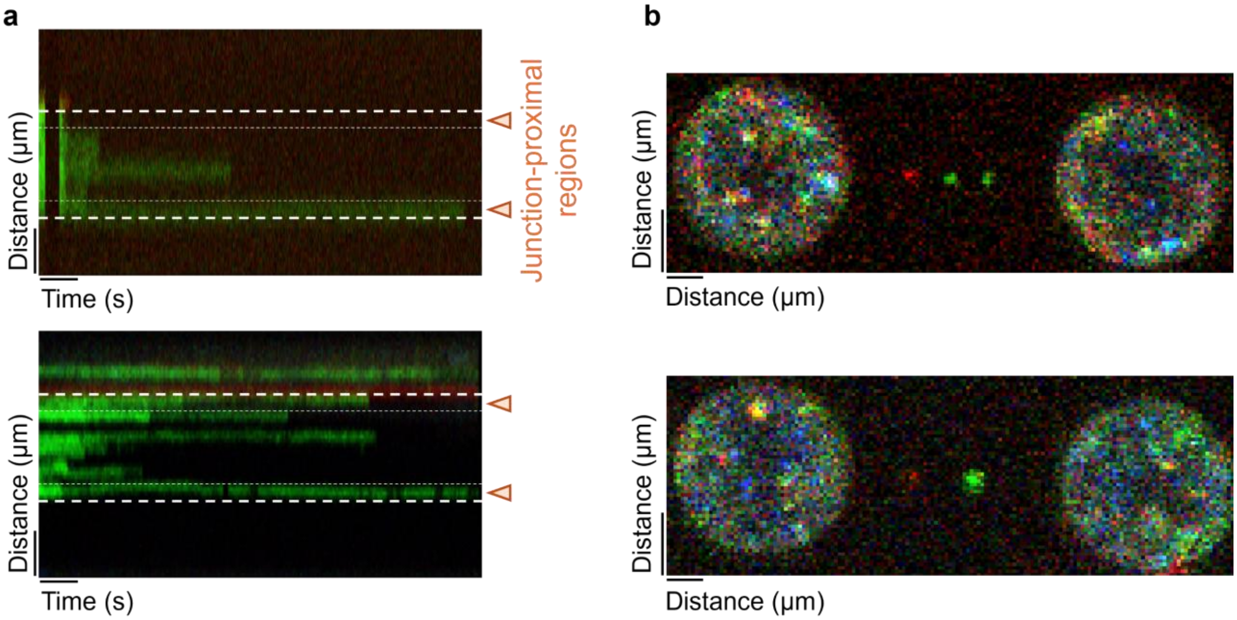


**Supplementary Figure. 3 | Facilitated exchange and initial engagement with ScRPA.**

**a**, Representative kymographs depicting facilitated exchange with fluorescently labeled ScRPA. **b**, Representative image scans depicting ScRPA signal (in green) as it initiates binding. The red label represents an ATTO647N dye.

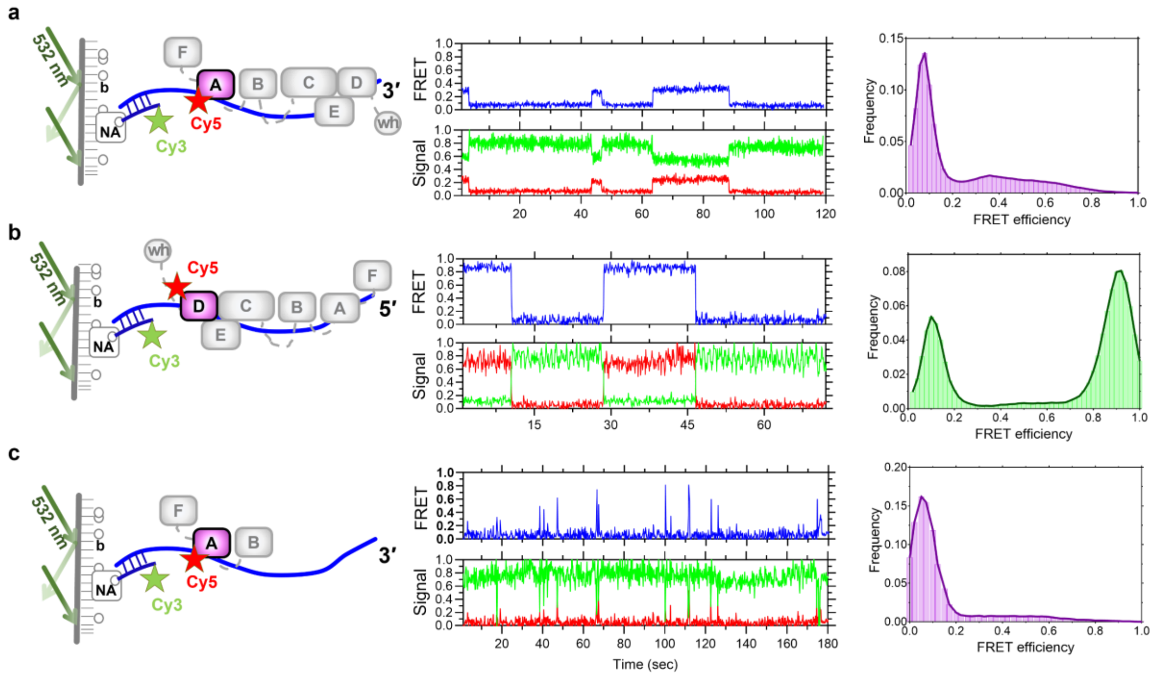


**Supplementary Figure 4 | Differences in dynamic DBA-A, and DBD-B binding relative to full-length RPA .**

**a**, TIRF setup to measure smFRET efficiency for RPA DBD-A$^{Cy5}$ and a Cy3-labeled 5' junction containing $(dT)_{30}$ overhang. The FRET efficiencies and signal anti-correlations point at a low-to-mid FRET state distribution. **b**, TIRF setup to measure smFRET efficiency for RPA DBD-D$^{Cy5}$ and a Cy3-labeled 3' junction containing $(dT)_{30}$ overhang. The FRET efficiencies and signal anti-correlations point at a long-lived, bimodal FRET states, moving between a low-to-high distribution. **c**, TIRF setup to measure smFRET efficiency for RPA DBD-D$^{Cy5}$ in an RPA-FAB construct, and a Cy3-labeled 5' junction containing $(dT)_{30}$ overhang. The FRET efficiencies and signal anti-correlations point at the highly dynamic binding behavior of the RPA-FAB domains.

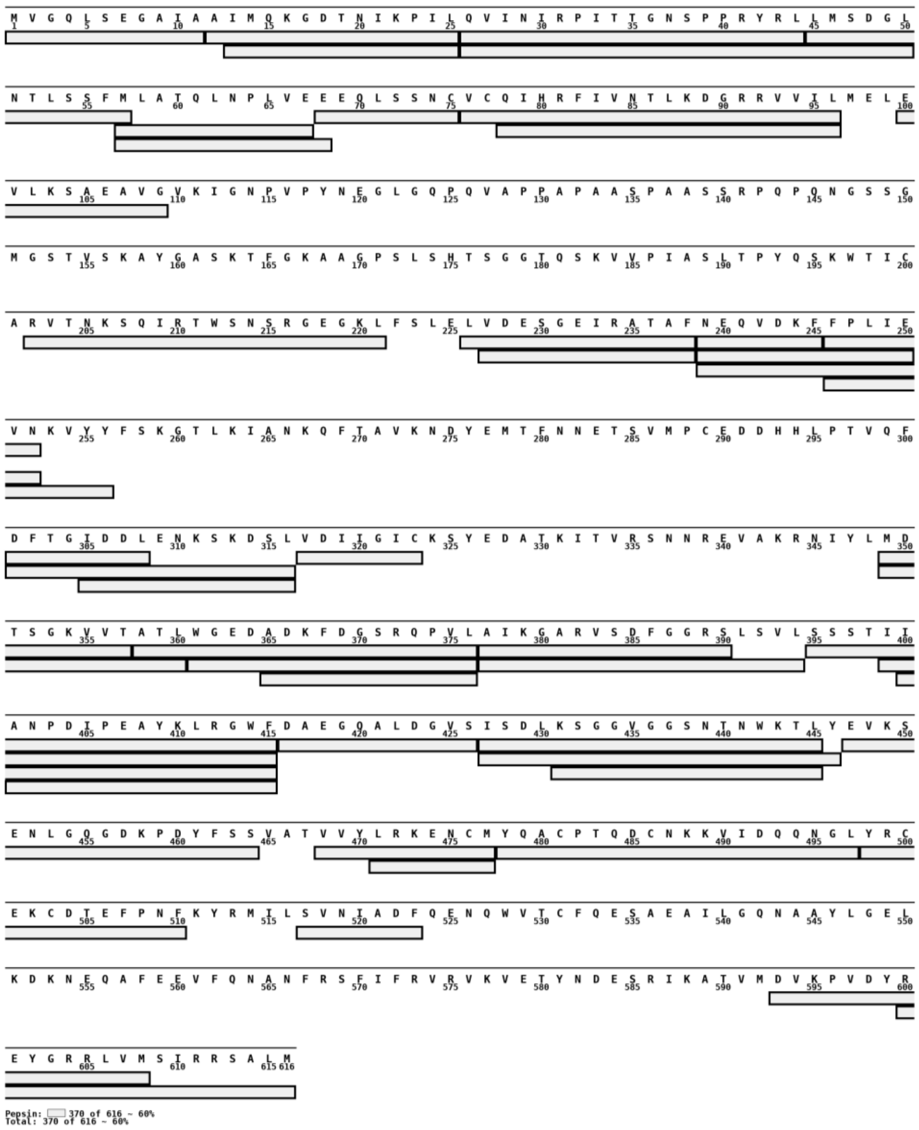

**Supplementary Figure 5 | Peptide coverage map of RPA 70.**

HDX-MS analysis resulted in a total of 47 peptides that were deemed highly confident, producing ~60% protein sequence coverage. Graphic was made using MSTools online software suite (Kavan and Man, 2010).

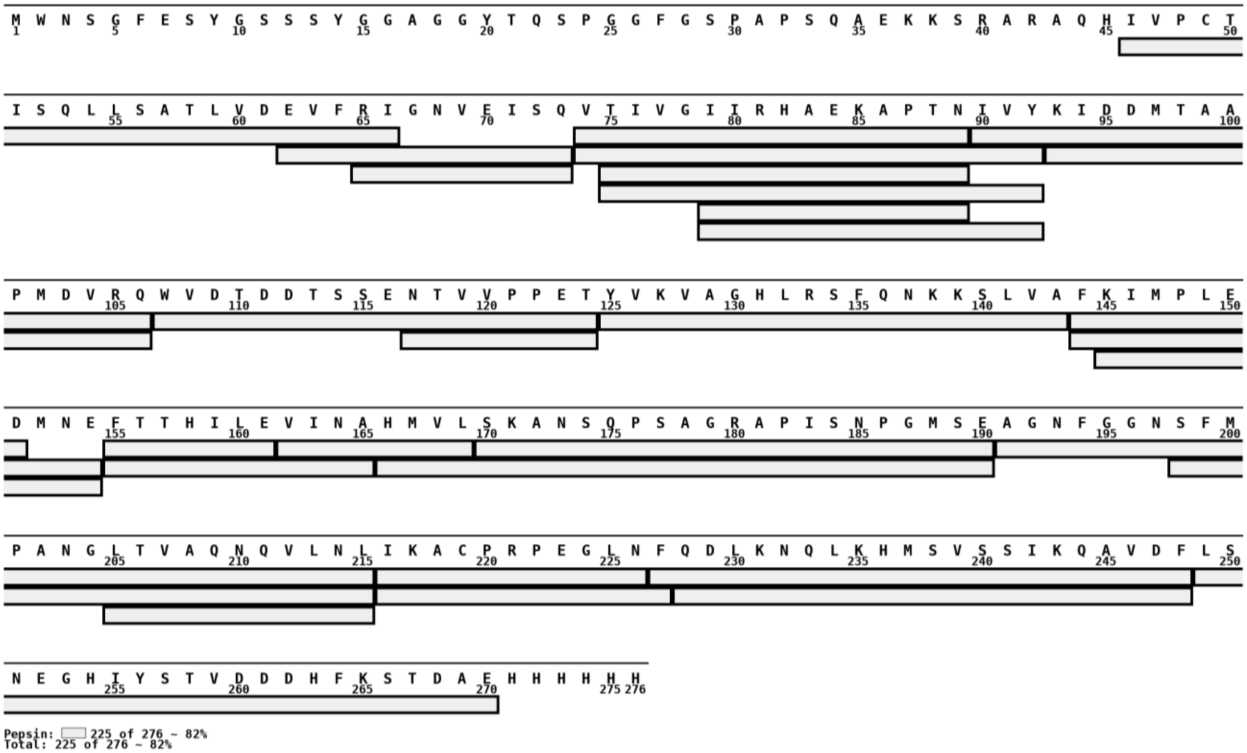


**Supplementary Figure 6 | Peptide coverage map of RPA 32.**

HDX-MS analysis resulted in a total of 30 peptides that were deemed highly confident, producing ~82% protein sequence coverage. Graphic was made using MSTools online software suite (Kavan and Man, 2010).

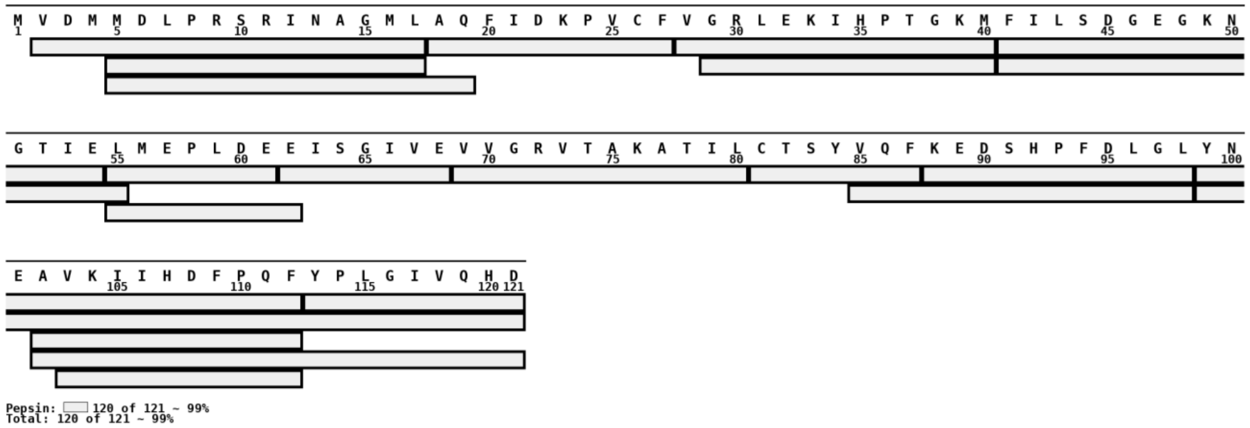


**Supplementary Figure 7 | Peptide coverage map of RPA 14.**

HDX-MS analysis resulted in a total of 21 peptides that were deemed highly confident, producing ~ 99% protein sequence coverage. Graphic was made using MSTools online software suite (Kavan and Man, 2010).

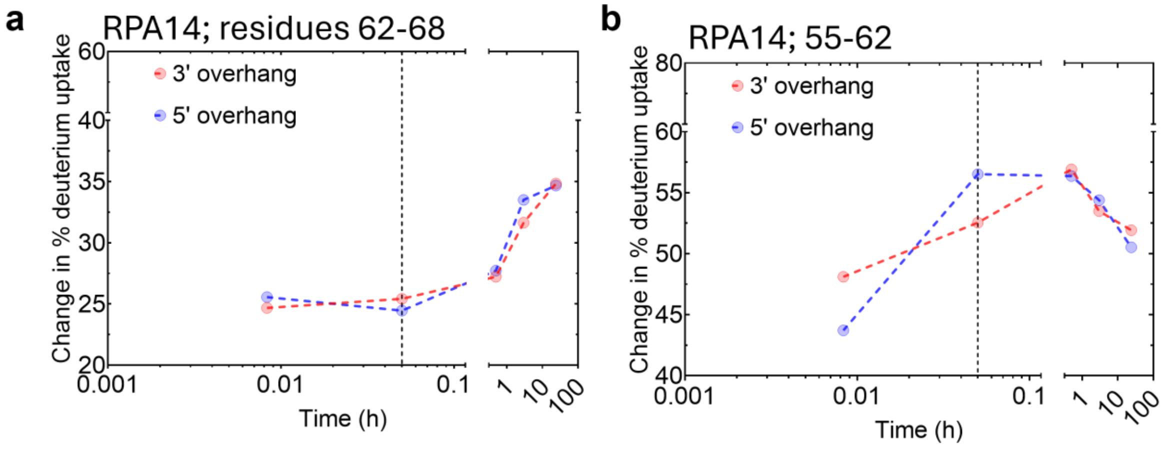


**Supplementary Figure 8 | Largest peptide-level change in percentage deuterium uptake for RPA14.**

**a-b**, Scatter-plots representing the change in percentage deuterium uptake as a function of time for the RPA14, regions spanning 62-68 (**a**), and from 55-62 (**b**). Dotted black line represents the three-minute time-point.

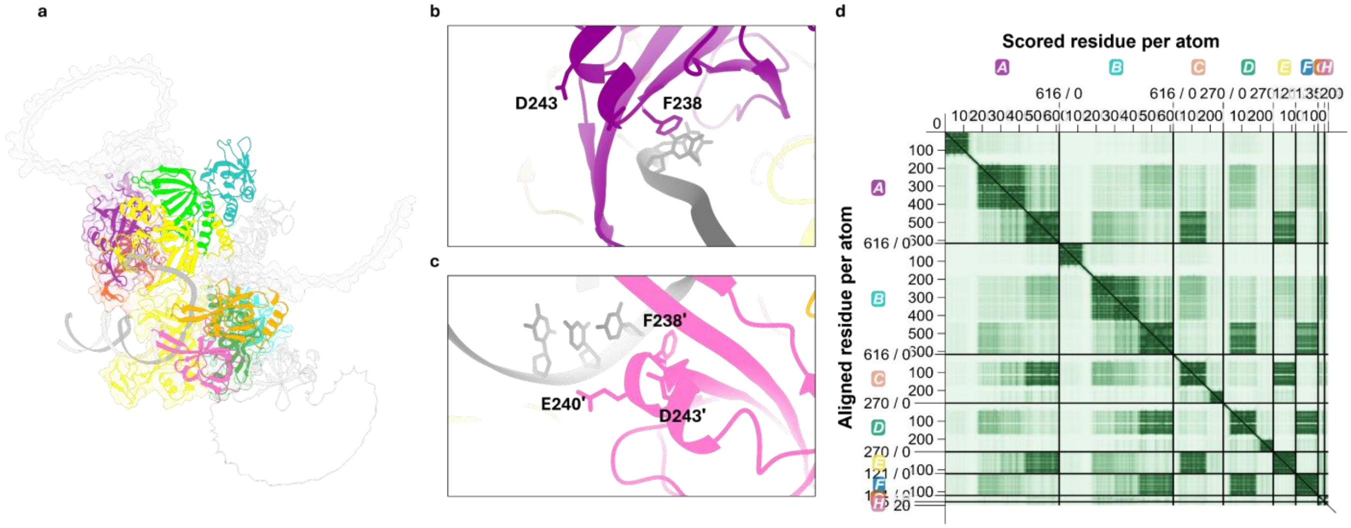


**Supplementary Fig. 9 | AlphaFold3 prediction of two RPA molecules bound to a 5′ junction (dT)15 ssDNA overhang.**

**a**, The composite AF3 model of both RPA molecules positioned on the junction-containing overhang substrate. The domains are color-coded, where DBD-A is magenta, DBD-B is orange, DBD-C is yellow, DBD-D is green, and OB-E is cyan. A dark and light shade of the same color was used to indicate the same OB-domain but in two different RPA molecules. substrate. **b and c**, insets representing the junction-specific motif, specifically F238, E240, and D243 within RPA DBD-A of both RPA70 subunits. **d**, the predicted alignment error (PAE) metric for the AF3 model depicting modest confidence in the relative positions of the motifs.

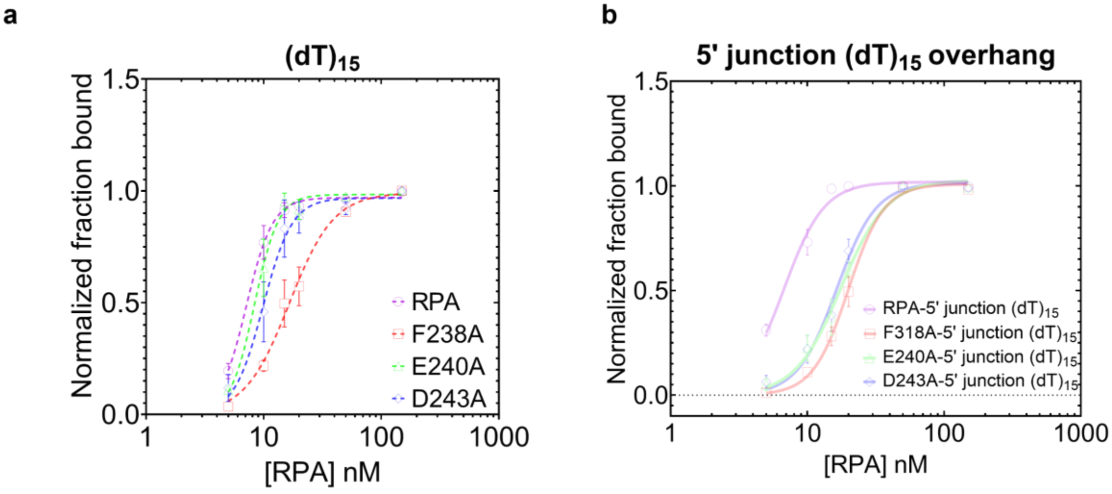


**Supplementary Figure. 10 | Binding properties of junction-specific mutants on various 15 nucleotide-long ssDNA substrates.**

**a**, Quantification of DNA binding (in terms of normalized fraction bound) for wild-type and mutant conditions of hRPA to a $(dT)_{15}$ ssDNA substrate. Data represents mean ± SEM from N= 3 repeats, and the dotted lines represent a non-linear specific binding fit with Hill slope model. **b**, Quantitation of DNA binding (in terms of normalized fraction bound) for wild-type and mutant conditions of hRPA to a 5′ junction $(dT)_{15}$ overhang. Data represents mean ± SEM from N ≥ 3 repeats, and the dotted lines represent the non-linear specific binding fit with Hill slope model.

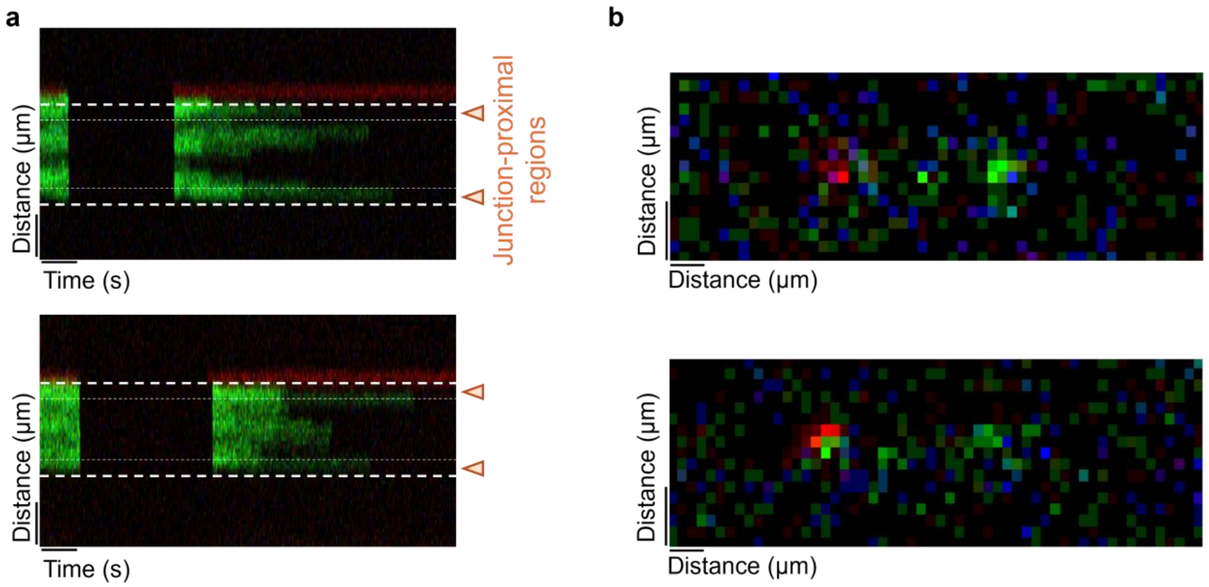


**Supplementary Figure. 11 | Facilitated exchange and initial engagement with hRPA-F238A.**

**a**, Representative kymographs depicting facilitated exchange with fluorescently labeled hRPA-F238A. **b**, Representative image scans depicting hRPA-F238A signal (in green) as it initiates binding. The red label represents an ATTO647N dye.

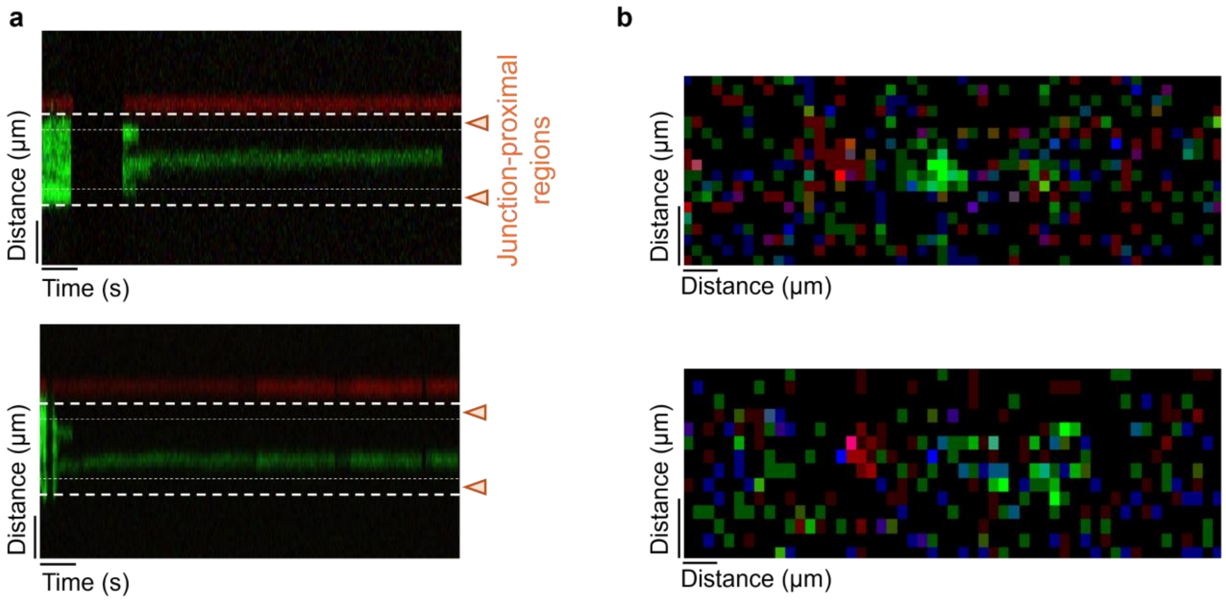


**Supplementary Figure. 12 | Facilitated exchange and initial engagement with hRPA-E240A.**

**a**, Representative kymographs depicting facilitated exchange with fluorescently labeled hRPA-E240A. **b**, Representative image scans depicting hRPA-E240A signal (in green) as it initiates binding. The red label represents an ATTO647N dye.

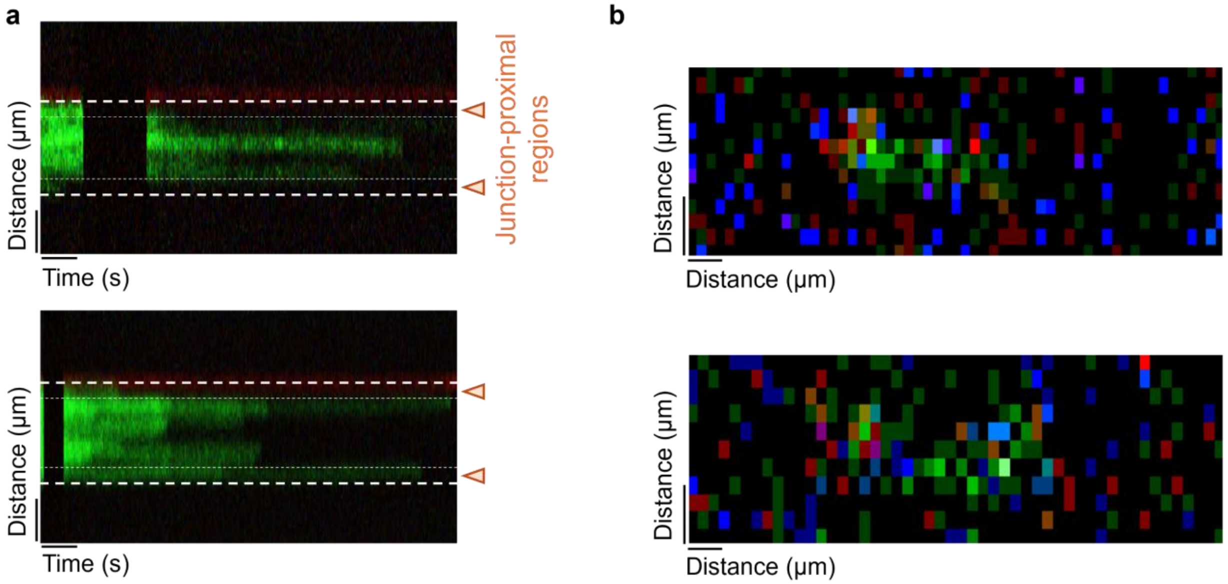


**Supplementary Figure. 13 | Facilitated exchange and initial engagement with hRPA-D243A.**

**a**, Representative kymographs depicting facilitated exchange with fluorescently labeled hRPA-D243A. **b**, Representative image scans depicting hRPA-D243A signal (in green) as it initiates binding. The red label represents an ATTO647N dye.

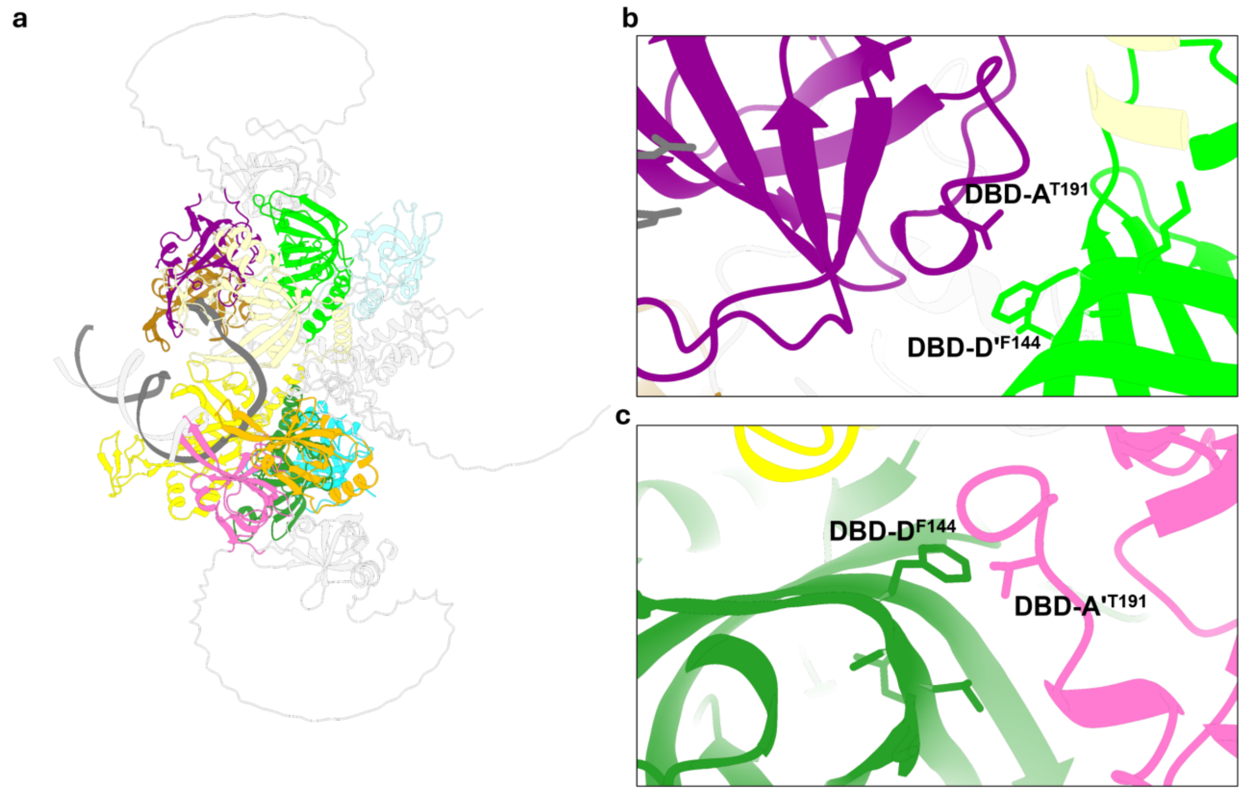


**Supplementary Figure 14 | AlphaFold3 model of two-bound RPA on a 5′ junction $(dT)_{15}$ overhang highlighting Thr191 in RPA70.**

**a**, The composite AF3 model of two RPA molecules bound to a 5′ junction-carrying $(dT)_{15}$ overhang substrate. **b and c**, insets representing the phosphorylation site – T191 within RPA DBD-A of both RPA70 subunits, and it's relative proximity to F144 of the adjacent RPA32 DBD-D in each case.

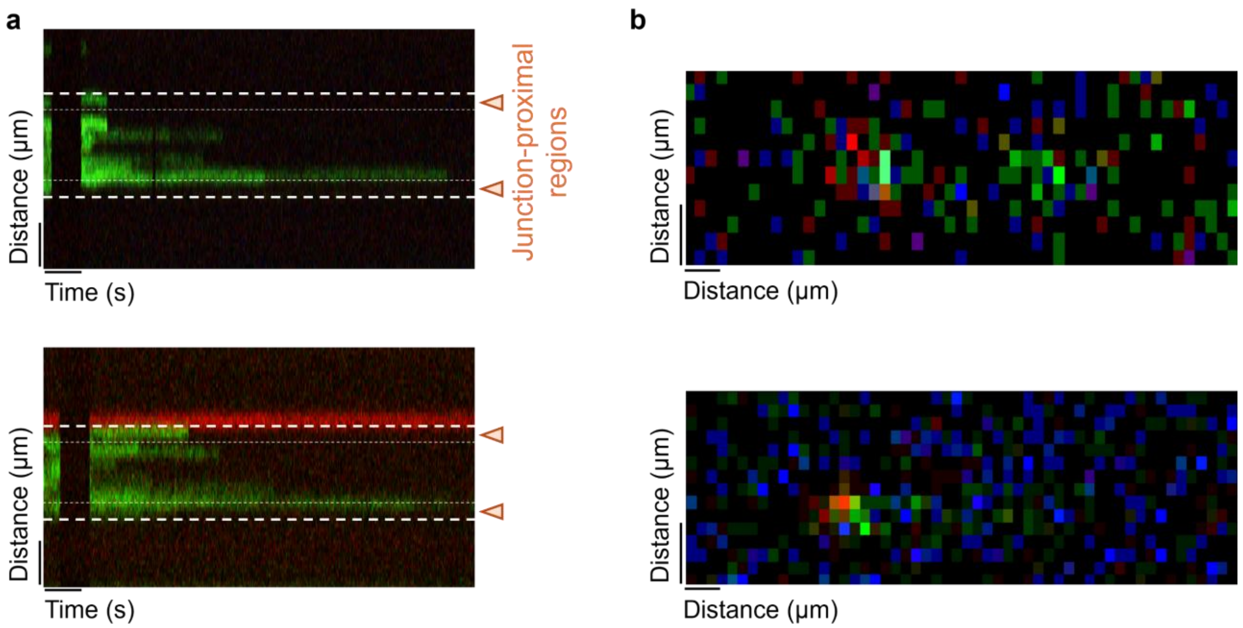


**Supplementary Figure. 15 | Facilitated exchange and initial engagement with hRPA-2D.**

**a**, Representative kymographs depicting facilitated exchange with fluorescently labeled hRPA-2D. **b**, Representative image scans depicting hRPA-2D signal (in green) as it initiates binding. The red label represents an ATTO647N dye.

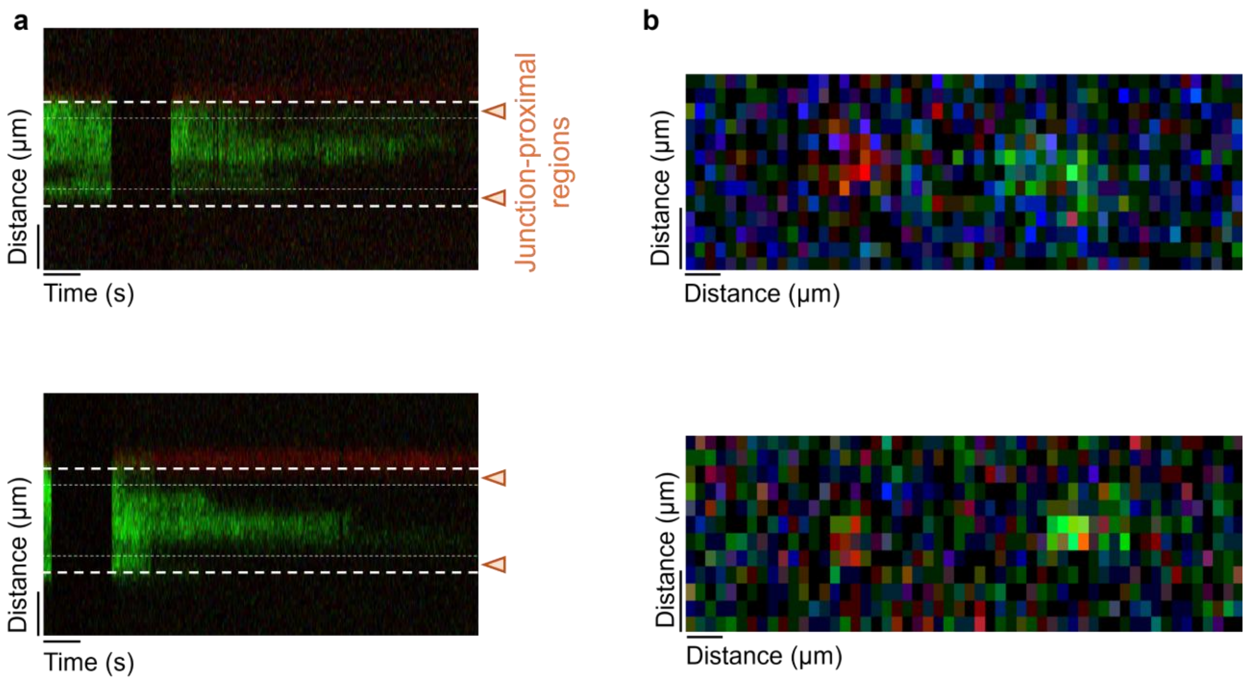


**Supplementary Figure. 16 | Facilitated exchange and initial engagement with hRPA-3D.**

**a**, Representative kymographs depicting facilitated exchange with fluorescently labeled hRPA-3D. **b,** Representative image scans depicting hRPA-3D signal (in green) as it initiates binding. The red label represents an ATTO647N dye.

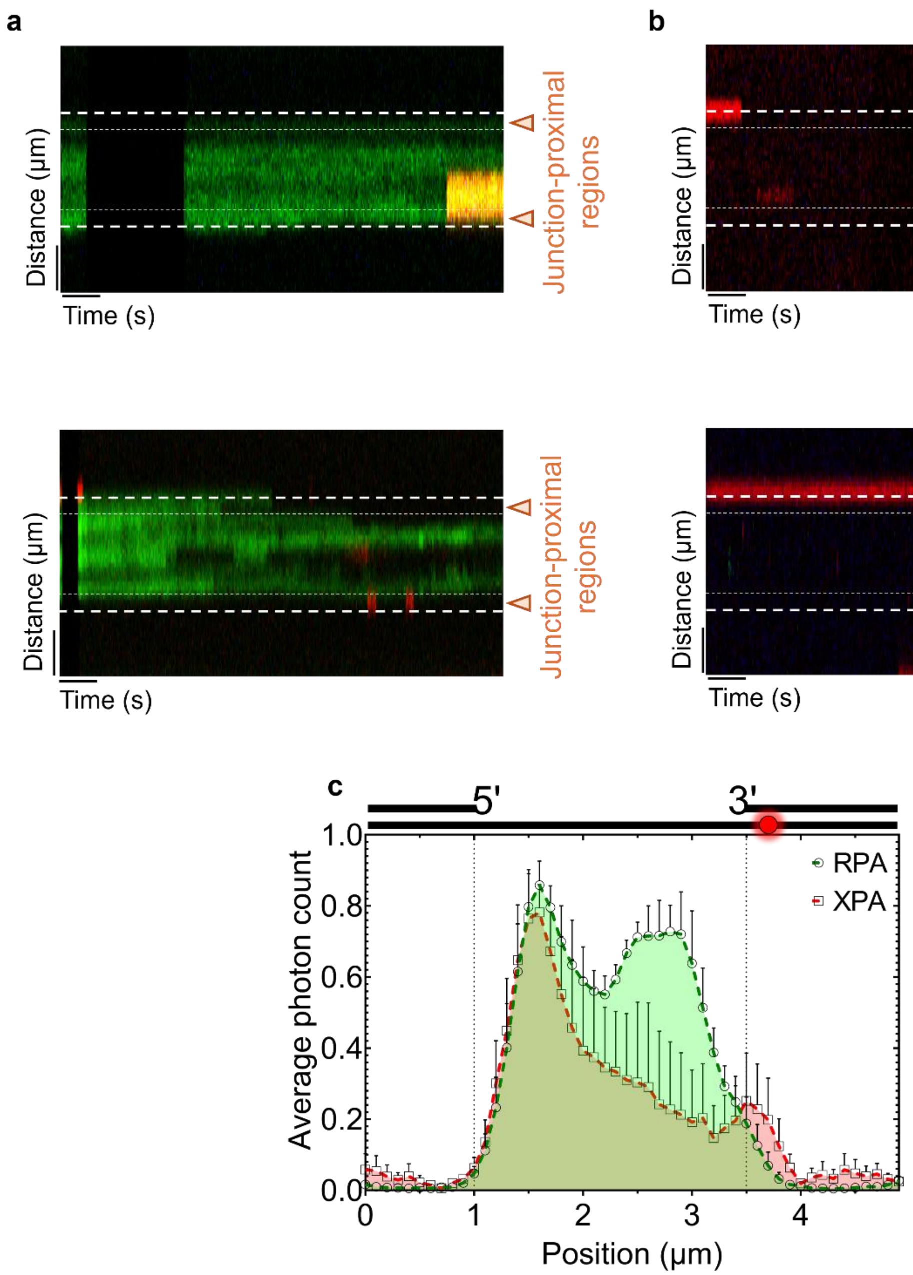


**Supplementary Figure. 17 | XPA$^{Cy5}$ binding to RPA-coated and non-RPA-coated gapped DNA substrates.**

**a,** Representative kymographs depicting XPA$^{Cy5}$ binding to RPA$^{MB543}$-coated gapped DNA substrate. **b**, representative kymographs depicting XPA$^{Cy5}$ encountering a gapped DNA substrate with no RPA$^{MB543}$.

**Supplementary Table 1 | Composition of the DNA oligonucleotides used in the study.**

| | |
|---|---|
| 35mer ssDNA for the 3′ junction $(dT)_{15}$ overhang | 5′ - TTT TTT TTT TTT TTT GCC TCG CTG CCG TCG CCA GG - 3′ |
| Complementary 20mer strand for the 3′ junction 35mer | 5′ - CCT GGC GAC GGC AGC GAG GC - 3′ |
| 35mer ssDNA for the 5′ junction $(dT)_{15}$ overhang | 5′ - GGA CCG CTG CCG TCG CTC CGT TTT TTT TTT TTT TT - 3′ |
| Complementary 20mer strand for the 5′ junction 35mer | 5′ - CGG AGC GAC GGC AGC GGT CC - 3′ |
| $(dT)_{15}$ | 5′ - TTTTTTTTTTTTTTT - 3′ |

**Supplementary Table 2 | AlphaFold3 input conditions used in this study.**

| | |
|---|---|
| RPA70 | MVGQLSEGAIAAIMQKGDTNIKPILQVINIRPITTGNSPPRYRLLMSDGLNTLSSFMLATQLNPLVEEEQL SSNCVCQIHRFIVNTLKDGRRVVILMELEVLKSAEAVGVKIGNPVPYNEGLGQPQVAPPAPAASPAASS RPQPQNGSSGMGSTVSKAYGASKTFGKAAGPSLSHTSGGTQSKVVPIASLTPYQSKWTICARVTNKSQI RTWSNSRGEGKLFSLELVDESGEIRATAFNEQVDKFFPLIEVNKVYYFSKGTLKIANKQFTAVKNDYEM TFNNETSVMPCEDDHHLPTVQFDFTGIDDLENKSKDSLVDIIGICKSYEDATKITVRSNNREVAKRNIYL MDTSGKVVTATLWGEDADKFDGSRQPVLAIKGARVSDFGGRSLSVLSSSTIIANPDIPEAYKLRGWFDA EGQALDGVSISDLKSGGVGGSNTNWKTLYEVKSENLGQGDKPDYFSSVATVVYLRKENCMYQACPTQ DCNKKVIDQQNGLYRCEKCDTEFPNFKYRMILSVNIADFQENQWVTCFQESAEAILGQNAAYLGELKD KNEQAFEEVFQNANFRSFIFRVRVKVETYNDESRIKATVMDVKPVDYREYGRRLVMSIRRSALM |
| RPA32 | MWNSGFESYGSSSYGGAGGYTQSPGGFGSPAPSQAEKKSRARAQHIVPCTISQLLSATLVDEVFRIGNV EISQVTIVGIIRHAEKAPTNIVYKIDDMTAAPMDVRQWVDTDDTSSENTVVPPETYVKVAGHLRSFQNK KSLVAFKIMPLEDMNEFTTHILEVINAHMVLSKANSQPSAGRAPISNPGMSEAGNFGGNSFMPANGLTV AQNQVLNLIKACPRPEGLNFQDLKNQLKHMSVSSIKQAVDFLSNEGHIYSTVDDDHFKSTDAE |
| RPA14 | MVDMMDLPRSRINAGMLAQFIDKPVCFVGRLEKIHPTGKMFILSDGEGKNGTIELMEPLDEEISGIVEV VGRVTAKATILCTSYVQFKEDSHPFDLGLYNEAVKIIHDFPQFYPLGIVQHD |
| ssDNA-1 | CCTGGCGACGGCAGGCTCCG |
| ssDNA-2 | TTTTTTTTTTTTTTTTTTTTTTTTTTTTTTGCCTCGCTGCCGTCGCCAGG |
| ssDNA-3 | GGACCGCTGCCGTCGCTCCGTTTTTTTTTTTTTTT |
| ssDNA-4 | CGGAGCGACGGCAGCGGTCC |

ssDNA 1 & 2 used to form the 3' junction $(dT)_{15}$ with 6 base pairs melted.

ssDNA 3 and 4 used to form the 5' junction $(dT)_{15}$.
